# The *Aspergillus fumigatus* C2-Domain Protein SppA is required for septal integrity and alters susceptibility to echinocandins and neutrophil killing during infection

**DOI:** 10.64898/2026.07.17.739181

**Authors:** Dante G Calise, Madeline L Michaelis, Jin Woo Bok, Zhili Chen, Joshua J Coon, Anna Huttenlocher, Benjamin J Chadwick, Nancy P Keller

## Abstract

*Aspergillus fumigatus* is a major opportunistic fungal pathogen whose ability to maintain hyphal integrity and withstand host defenses is critical for virulence. Septal pores, which connect hyphal compartments, are dynamically regulated to preserve cellular integrity under stress, yet the molecular components governing this process remain incompletely defined. Here, we identify and characterize a septal pore-associated protein, SppA, and demonstrate its essential role in maintaining septal integrity in *A. fumigatus*. We show that expression of SppA is positively regulated by the transcription factor ZfpA and is induced in response to the cell wall-targeting antifungal caspofungin. Deletion of *sppA* resulted in defective septal organization and increased susceptibility to hyphal damage. The mutant exhibited heightened sensitivity to cell wall-targeting antifungal agents, indicating a role in cell wall stress tolerance. In a zebrafish model of invasive aspergillosis, loss of SppA significantly attenuated virulence which was abrogated in neutrophil-deficient zebrafish. Further, the mutant strain displayed increased susceptibility to killing by primary human neutrophils, suggesting that proper septal pore formation contributes to fungal survival during host immune attack. Together, our findings establish SppA as a critical determinant of septal integrity, antifungal tolerance, and pathogenicity in *A. fumigatus*, and position it as part of a ZfpA-regulated, caspofungin-responsive pathway that supports fungal survival during stress and infection.

**Author Summary:** *Aspergillus fumigatus* is a common environmental mold that can cause life-threatening infections in people with weakened immune systems. For successful invasion of host tissue, the fungus requires the ability to protection sections of its hyphae from cell wall targeting antifungals and host immune cell attack by closing septal (cross wall) pores distributed throughout hyphal strands. We have identified an *A. fumigatus* protein, SppA, required for proper septal pore closure. Loss of SppA reduces the ability of hyphae to withstand treatment with antifungals and the ability of *A. fumigatus* to cause disease in a zebrafish infection model. The SppA mutant was particularly susceptible to killing by neutrophils, key immune cells that help control fungal infections. Our findings reveal an important mechanism that helps *A. fumigatus* survive environmental and host-imposed stresses and highlight septal pore regulation as a potential target for future antifungal strategies.

## Introduction

Invasive fungal infections pose a significant threat to human health with extremely high mortality rates compared to other common infectious diseases [1]. Invasive aspergillosis (IA), most often caused by the ubiquitous environmental saprotroph *Aspergillus fumigatus*, is estimated to claim more than one million lives annually. One of the greatest challenges presented by invasive fungal infections is a severely limited arsenal of antifungals used in the treatment of these pathogens. Currently only one class of drug—the triazoles, targeting the cell membrane—is approved for primary therapy of IA [2]. Fungi and animals share many homologous cellular processes which makes the identification of fungal-specific drug targets without host toxic effects particularly difficult as was observed with amphotericin B, the first approved cell membrane-targeting antifungal drug for IA [3,4]. Since animal cells lack a cell wall, the fungal cell wall presents an enticing target for antifungal drug identification. However, remarkable diversity in cell wall composition among human fungal pathogens has presented a major obstacle in identifying a “one size fits all” drug [5,6].

Caspofungin, micafungin, and anidulafungin comprise the echinocandin antifungal class of semisynthetic lipopeptides targeting the cell wall synthase β-1,3-glucan synthase (FksA) leading to reduction in β-1,3-glucans, key components of the fungal cell wall [7,8]. However, these drugs have varying efficacy against different fungal pathogens from highly fungicidal activity against the yeast pathogen *Candida albicans* to almost no activity against *Cryptococcus* and the coenocytic molds which cause mucormycosis [9]. In the case of *A. fumigatus* and other pathogenic aspergilli, the efficacy of these drugs is less well defined. Currently, echinocandins are only approved for treatment of IA in salvage therapy when azoles alone fail to control the infection. However, *in vitro,* echinocandins exhibit both fungistatic and limited fungicidal activity against *A. fumigatus,* incentivizing research to identify molecular targets to enhance their clinical efficacy in treating *Aspergillus* infections [10,11]. Recently, a new class of antifungal called fungerps was named following the approval of the semisynthetic terpene ibrexafungerp, derived from the fungal natural product enfumafungin, for treatment of vulvovaginal candidiasis. Although structurally distinct from the echinocandins, this compound also targets FksA, thus exhibiting a similar antifungal profile against fungal pathogens. Importantly, the much smaller molecular size of fungerps relative to echinocandins allows oral administration whereas the latter must be administered intravenously [12]. Ibrexafungerp is currently in late-stage clinical trials for use in IA treatment, as rising incidence of azole resistance among *A. fumigatus* isolates presents a growing demand for novel therapeutic approaches and the enhancement of already approved second line antifungals like the echinocandins and fungerps [13].

Inhibition of FksA in *A. fumigatus* disrupts proper cell wall synthesis at the growing tip leading to lysis of the apical compartment [10]. Despite this, the fungus can limit fungicidal damage by rapidly occluding septal pores with Woronin bodies, thereby restricting cytoplasmic loss. This response permits the formation of a new growth axis from the sealed septum and enables hyphal recovery [14]. Consistent with this, septation is essential for preventing complete killing of *A. fumigatus* hyphae by FksA inhibiting antifungals [15,16]. In parallel, *A. fumigatus* remodels its cell wall during sustained drug exposure, with rearrangements of chitin, chitosan, and α-1,3-glucan contributing to adaptive survival [17]. However, the regulatory mechanisms coordinating these protective processes remain incompletely defined. Notably, our previous work identified the transcription factor ZfpA as a central regulator of this adaptive response: ZfpA promotes chitin synthesis during echinocandin stress, increases septation, and enhances tolerance to the fungicidal activity of these drugs *in vitro* and during *in vivo* infection [18–21]. These findings position ZfpA as a critical node linking cell wall remodeling and septation-mediated survival during antifungal and host immune challenge.

Here, we performed an untargeted proteomic screen of *A. fumigatus* ZfpA deletion and overexpression mutants during caspofungin treatment to identify downstream effectors that mediate hyphal protection against FksA inhibition. This approach identified the C2-domain protein SppA as a strongly induced, ZfpA-dependent factor during echinocandin stress. Functional analyses revealed that SppA is required for tolerance not only to FksA-targeting antifungals but broadly across cell wall and osmotic stress conditions. Mechanistically, our data support a role for SppA in promoting chitin deposition and proper septal formation, processes that are essential for antifungal tolerance. Consistent with this, SppA is also required for full virulence during invasive infection in a neutrophil dependent manner. Together, these findings define SppA as a key downstream effector of ZfpA that links cell wall remodeling and septation to antifungal tolerance and pathogenicity.

## Results

### The C2 domain protein SppA is regulated by the transcription factor ZfpA during caspofungin treatment

The transcription factor ZfpA is required for tolerance to FksA-targeting antifungals [18–20]. To identify ZfpA regulated proteins mediating echinocandin tolerance, we performed a proteomic screen comparing *A. fumigatus* protein expression in wild type (WT), Δ*zfpA*, and OE::*zfpA* strains under caspofungin treatment (Supplemental Data 1). Confirming our previous work and providing confidence in our approach, we observed the increased expression of PpoA, known to be induced by ZfpA in response to caspofungin (Figure 1A) [18]. Interested in uncharacterized proteins, we noted AFUA_2G01790 was the most upregulated protein in OE::*zfpA* vs. Δ*zfpA* (Figure 1A) and the most downregulated in Δ*zfpA* vs. WT (Supplemental Figure 1) during caspofungin treatment. We identified AFUA_2G01790 as an orthologue of the recently characterized *Aspergillus oryzae* SppA, localized to the septal pore and required for septal integrity during hypotonic shock [22], which we hereafter refer to as *A. fumigatus* SppA. SppA abundance was ∼67-fold lower in Δ*zfpA* than WT during caspofungin treatment, with no significant difference under drug-free conditions (Supplemental Data 1). Consistent with these findings, Northern blot analysis showed increased *sppA* transcript levels in OE::*zfpA* compared to WT and Δ*zfpA*, which were indistinguishable without drug (Supplemental Figure 1). Together, these data demonstrate that SppA is induced by caspofungin in a ZfpA-dependent manner.

**Figure 1.** The C2 domain containing protein SppA is required for tolerance to β-1,3-glucan synthase (FksA) targeting antifungal drugs. **(A)** Log_2_FC abundance of *A. fumigatus* Af293 proteins in OE::*zfpA* relative to Δ*zfpA* extracted from mycelia after treatment with 0.5 µg/mL caspofungin for four hours. Points representing PpoA and SppA are colored yellow. SppA was identified as the most highly upregulated protein. **(B)** Percent survival of *A. fumigatus* Af293 germlings treated with 2 µg/mL caspofungin (CSF) after 16 hours in GMM + 0.01% yeast extract. ‘****’ denotes p < 0.0001 calculated by one-way ANOVA with Tukey’s multiple comparisons. **(C)** Dilution plating of *A. fumigatus* strains on glucose minimal media with caspofungin (CSF), enfumafungin (ENF), or DMSO vehicle control after incubation at 37°C for 48 hours. **(D)** Percent survival of *A. fumigatus* Af293 germlings treated with 2 µg/mL caspofungin after 16 hours in GMM + 0.01% yeast extract. Strains with p values less than 0.05 calculated by one-way ANOVA with Tukey’s multiple comparisons are indicated by distinct letters. **(E)** Dilution plating of *A. fumigatus* strains on glucose minimal media with CSF, micafungin (MCF), ENF, or DMSO vehicle control after incubation at 37°C for 48 hours.

### SppA is required for tolerance to FksA targeting antifungals

To assess the role of SppA in tolerance to FksA targeting antifungals, we generated a deletion mutant (Δ*sppA*), complemented strain (*sppA*+) and overexpression strain (OE::*sppA*) in the *A. fumigatus* 293 genetic background (Supplemental Figure 2). Germling survival following 16 hours of treatment with 2 µg/mL caspofungin was then measured using previously published methods [18]. We found that loss of SppA rendered germlings highly susceptible to fungicidal killing by caspofungin, and this sensitivity was abolished in the complemented strain. The OE::*sppA* mutant was indistinguishable to the WT strain, suggesting that wild type expression is sufficient for maximal protection against caspofungin mediated tip lysis (Figure 1B). Furthermore, *sppA* deletion and complement strains in the *A. fumigatus* CEA10 lineage (Supplemental Figure 2) yielded similar results as determined by caspofungin and enfumafungin treatments, thus demonstrating a conserved role for SppA in tolerance to FksA inhibiting drugs (Figure 1C).

To determine whether SppA functions downstream of ZfpA in antifungal tolerance, we generated Δ*sppA* and OE::*sppA* mutants in the OE::*zfpA* and Δ*zfpA* backgrounds, respectively (Supplemental Figure 2). Consistent with previous observations, OE::*zfpA* germlings exhibited enhanced survival relative to WT. However, deletion of *sppA* in the *OE::zfpA* background abolished this phenotype, resulting in 100% germling killing by 16 hours due to caspofungin-induced tip lysis (Figure 1D). The OE::*zfpA* Δ*sppA* double mutant also demonstrated drastic sensitivity to caspofungin, micafungin, and enfumafungin on solid media (Figure 1E). Interestingly, overexpression of *sppA* did not protect Δ*zfpA* germlings from caspofungin-induced killing suggesting that SppA alone is insufficient to confer tolerance and likely acts in concert with other ZfpA regulated factors (Figure 1D). Consistent with this finding, the Δ*zfpA* OE::*sppA* double mutant did not exhibit appreciably improved growth relative to the Δ*zfpA* strain on solid media (Supplemental Figure 3). Taken together, these results indicate that ZfpA dependent activation of SppA is necessary, but not sufficient, for tolerance to FksA targeting antifungals and that SppA functions alongside other ZfpA regulated proteins to promote echinocandin and enfumafungin tolerance.

### SppA directs chitin synthesis necessary for proper septal formation

Since ectopically expressed SppA-GFP localizes to septa of *A. oryzae* [22], we stained WT, Δ*sppA*, *sppA*+, and OE::*sppA* hyphae with calcofluor white and performed high resolution fluorescent microscopy to visualize septal structures. Compared to WT and *sppA+* strains, Δ*sppA* hyphae consistently demonstrated poorly formed or incomplete septa (Figure 2A). Although OE::*sppA* formed complete septa, it also displayed intensely calcofluor white stained cell wall patches, suggesting aberrant chitin deposition associated with *sppA* overexpression (Figure 2A). Despite these morphological changes, the mean pixel intensity of calcofluor white staining did not differ in Δ*sppA* or OE::*sppA* mutant compared to WT suggesting total hyphal chitin/chitosan content was not substantially altered (Supplemental Figure 4). Notably, chitin-rich cell wall patches were still observed in the Δ*zfpA* OE::*sppA* double mutant; however, the reduced septation phenotype associated with Δ*zfpA* was not restored, suggesting that SppA-mediated chitin deposition and septum initiation are at least partially separable processes (Figure 2B and Supplemental Figure 4). Interestingly, deletion of *sppA* in the OE::*zfpA* background resulted in both malformed septa and brightly stained chitin patches in the cell wall, further supporting the hypothesis that other ZfpA-regulated proteins interact with SppA to coordinate chitin synthesis in septal formation (Figure 2C).

**Figure 2.** SppA is indispensable for proper septal formation. **(A)** Representative micrographs of *A. fumigatus* Af293 WT, Δ*sppA*, *sppA*+, and OE::*sppA* hyphae stained with calcofluor white before fluorescent imaging. **(B)** Representative micrographs of *A. fumigatus* Af293 Δ*zfpA* and Δ*zfpA* OE::*sppA* hyphae stained with calcofluor white before fluorescent imaging. **(C)** Representative micrographs of *A. fumigatus* Af293 OE::*zfpA* and OE::*zfpA* Δ*sppA* hyphae stained with calcofluor white before fluorescent imaging. **(A-C)** Insets show septa digitally enlarged by 2X. Red arrowheads indicated brightly stained cell wall patches consistently observed in the OE::*sppA* mutants as well as in the OE::*zfpA* Δ*sppA* strain. Scale bars represent 20 µM.

### SppA-GFP forms contracting rings during septation

In order to determine the subcellular localization of SppA in *A. fumigatus* we tagged the C-terminus of SppA with GFP in both a WT and OE::*zfpA* background (Supplemental Figure 5). Confocal imaging revealed that SppA-GFP assembled into contractile ring-like structures during septum formation (Figure 3A-B). To assess whether SppA contributes to the ectopic chitin patches observed in *OE:sppA* hyphae, we overexpressed SppA-GFP under the *A. nidulans gpdA* promoter (Supplemental Figure 5). Overexpressed SppA-GFP localized diffusely throughout the cytoplasm but was enriched at septal rings and hyphal tips, suggesting roles in both septation and apical growth (Figure 3C). Additionally, bright GFP puncta colocalized with CFW stained chitin patches in the cell wall, supporting the role for SppA in directing chitin synthase activity (Supplemental Figure 6).

**Figure 3.** SppA-GFP localizes to sites of active septation and growing hyphal tips. **(A)** Representative confocal micrograph of OE::*zfpA sppA-GFP* hypha grown for 13 hours at 37° C in glucose minimal media. Yellow arrowhead indicates an early forming septum, cyan arrowhead indicates a late forming septum, and red arrowhead indicates a fully formed septum. Scale bar represents 10 µm. Insets are digitally enlarged by 2.5X. **(B)** Representative confocal images of an OE::*zfpA sppA-GFP* hypha grown for 13 hours at 37° C in glucose minimal media imaged every 2 min for 10 minutes. Scale bars represent 5 µm. **(C)** Representative confocal micrograph of OE::*sppA-GFP* hypha grown for 13 hours at 37° C in glucose minimal media. Yellow arrowhead indicates an early forming septum, cyan arrowhead indicates a late forming septum, and red arrowhead indicates a fully formed septum. Scale bar represents 10 µm. Insets are digitally enlarged by 2.5X. **(A-C)** Images in the GFP channel show a maximum intensity projection of Z-stacks imaged at a 0.2-µm interval with the DIC channel images collected in the center most plane of the Z-stack.

### SppA confers tolerance to cell wall and osmotic stress

Considering that septal pore plugging was compromised in an *A. oryzae* Δ*sppA* mutant [22], we deleted *sppA* in an *A. fumigatus* CEA10 strain constitutively expressing the fluorescent protein mMaroon1 (mM1) to observe cytoplasmic loss upon tip lysis under caspofungin treatment (Supplemental Figure 2). Whereas tip lysis resulted in loss of fluorescent signal only up to the apical most septum in the WT strain, tip lysis caused a complete loss of signal in the entire hyphae of the Δ*sppA* strain indicating a defect in septal pore plugging (Figure 4A). Since septal integrity is broadly important in the tolerance of cell wall and osmotic stress, we assessed the impact of *sppA* deletion on the susceptibility to physiological stressors in both WT and OE::*zfpA* backgrounds (Figure 4B). The Δ*sppA* mutants exhibited increased sensitivity to the cell wall perturbing agents Congo red and calcofluor white, as well as to hypertonic stress imposed by sorbitol or NaCl, relative to their corresponding WT and OE::*zfpA* controls (Figure 4B). Conversely, the mutants showed little-to-no difference in susceptibility to the membrane targeting antifungals voriconazole and amphotericin B (Figure 4C). Together, these data indicate that loss of SppA disrupts septal architecture and pore-plugging function, preventing compartmentalization following cell wall rupture due to overwhelming turgor pressure at the growing tip and leading to extensive cytoplasmic loss.

**Figure 4.** SppA is required for protective septal pore plugging in response to cell wall and osmotic stress. **(A)** Representative micrographs of WT and Δ*sppA A. fumigatus* CEA10 expressing cytosolic mMaroon1 (magenta) during growth in GMM with 2 µg/mL CSF imaged for 18 hours by brightfield and fluorescent microscopy. Visible septa at the final timepoint are indicated with black arrowheads. Scale bar represents 50 µm. **(B)** Dilution plating of *A. fumigatus Af293* strains on glucose minimal media with inhibitory levels of Congo red (CR) calcofluor white (CFW), Sorbitol, or NaCl after incubation at 37°C for 48 hours. **(C)** Dilution plating of *A. fumigatus* Af293 strains on glucose minimal media with voriconazole (VOR), amphotericin B (AmB), or DMSO vehicle control after incubation at 37°C for 48 hours.

### Loss of SppA severely attenuates virulence during invasive infection of larval zebrafish

Since aseptate mutants of *A. fumigatus* fail to demonstrate tissue invasive growth in a murine model of IA [15], we hypothesized that the septal defects observed in the Δ*sppA* mutant would similarly impair fungal virulence. To assess the impact of SppA on virulence, we utilized the established larval zebrafish model of IA [23]. Larval zebrafish were microinjected two days post fertilization with approximately forty *A. fumigatus* conidia or a mock PBS injection into the hindbrain ventricle to assess survival out to seven days under dexamethasone treatment to broadly suppress the host immune response (Figure 5A-B). In a dexamethasone treated model, larvae infected with *A. fumigatus* Af293 Δ*sppA* were nearly sevenfold less likely to die than those infected with the WT strain as determined by the Cox proportional-hazard model (Figure 5A). Similarly, loss of SppA in the *A. fumigatus* CEA10 background markedly attenuated virulence while virulence was restored to WT levels by ectopic complementation of *sppA* (Figure 5B). Notably, survival of larvae injected with CEA10 Δ*sppA* was not statistically distinguishable from PBS injected control indicating that SppA nearly abolishes pathogencity in this model (Figure 5B).

**Figure 5.** Loss of SppA drastically attenuates *A. fumigatus* virulence during invasive infection of larval zebrafish. **(A)** Pooled survival of WT zebrafish larvae injected with *A. fumigatus* Af293 and treated with 10 µM dexamethasone (DXM) out to seven days post injection (DPI) from five independent experiments. **(B)** Pooled survival of WT larvae injected with *A. fumigatus* CEA10 and treated with 10 µM DXM out to seven days post infection from three independent experiments. **(C)** Pooled survival of WT zebrafish larvae injected with *A. fumigatus* CEA10 and treated with 10 µM DXM for five days from three independent experiments. **(A-C)** *D. rerio* larvae were microinjected with ∼40 *A. fumigatus* conidia or mock PBS control into the hindbrain ventricle two days post fertilization **(A,B)** or the developing swim bladder four days post fertilization **(C)**. Eighteen to thirty fish were included for each condition per experimental replicate with four fish homogenized immediately following injection and plated to confirm similar injection CFUs between fungal strains. Pairwise Cox proportional hazard ratios were calculated with Sidak’s correction for multiple comparisons.

Since the hindbrain ventricle is only transiently observed during zebrafish development, infection of the swim bladder, a permanent air-filled compartment, has been proposed as a more physiologically relevant model of pulmonary infection. To test the role of SppA in this setting, we injected *A. fumigatus* CEA10 conidia (WT, Δ*sppA*, or *sppA*+) or PBS alone as a mock control, into the developing swim bladders of larvae four days post fertilization and monitored survival out to five days under dexamethasone treatment (Figure 5C). Consistent with the hindbrain infection model, larvae infected with the Δ*sppA* strain exhibited increased survival to those infected with the WT and complement strains (Figure 5C).

### SppA is required for tolerance to antifungal activity by echinocandins and neutrophils *in vivo*

Finally, since neutrophils are the key immune cell involved in hyphal killing, we utilized the neutrophil deficient zebrafish line overexpressing the dominant negative Rac2^D57N^ mutation under the neutrophil specific mpx promoter to assess the impact of SppA on echinocandin tolerance during *in vivo* infection [19]. Two days post fertilization, Tg(mpx:mCherry-2A-Rac2^D57N^) larvae were microinjected with *A. fumigatus* CEA10 WT, Δ*sppA*, *sppA*+, or mock PBS into the hindbrain ventricle and treated with micafungin or a vehicle control tracking survival out to seven days (Figure 6A). In contrast to infection of control (mpx:Rac2^WT^) larvae, hindbrain infection with the CEA10 Δ*sppA* mutant drastically reduced survival of neutrophil deficient (mpx:Rac2^D57N^) larvae compared to a mock PBS injection (Figure 6A-B). However, neutrophil deficient larvae were still significantly more likely to succumb to infection by the WT and complement strains compared to the Δ*sppA* mutant. This suggests an important role for SppA in resistance to neutrophil killing during infection, but it does not preclude the contribution of other factors to the reduced virulence of the Δ*sppA* mutant. Treatment with micafungin uniquely abolished virulence of the Δ*sppA* strain in neutrophil deficient larvae while it did not impact survival of larvae infected with the WT or complement strains in agreement with our *in vitro* results. This suggests that echinocandin treatment can compensate for a deficient neutrophil response to abolish the virulence of the Δ*sppA* mutant (Figure 6A).

**Figure 6.** SppA contributes to hyphal tolerance to fungicidal activity of host neutrophils and echinocandin drugs. **(A)** Pooled survival of mpx:mCherry-2A-Rac2^D57N^ transgenic zebrafish larvae injected with *A. fumigatus* CEA10 followed out to seven days with or without 0.5 µg/mL MCF treatment from three independent experiments. **(B)** Pooled survival of mpx:mCherry-2A-Rac2^WT^ transgenic zebrafish larvae injected with *A. fumigatus* CEA10 followed out to seven days from three independent experiments. **(A,B)** Zebrafish (*D. rerio*) larvae were microinjected with ∼40 *A. fumigatus* conidia or mock PBS control into the hindbrain ventricle two days post fertilization. Twenty to thirty fish were included for each condition per experimental replicate with four fish homogenized immediately following injection and plated to confirm similar injection CFUs between fungal strains. Pairwise Cox proportional hazard ratios were calculated with Sidak’s correction for multiple comparisons. **(C)** Probability of death of *A. fumigatus* CEA10 WT and Δ*sppA* hyphae due to *in vitro* killing by human PMNs isolated from three unique donors. Death curve represents three pooled experiments following fifty germlings of each strain per experiment tracked out to twelve hours. P < 0.0001 was determined by Logrank (Mantel-Cox) test. **(D)** Representative micrographs of fluorescent (OE::mMaroon1) WT and Δ*sppA A. fumigatus* hyphae co-incubated with primary human neutrophils 0, 6, and 12 hours after adding the neutrophils.

Next, we utilized the aforementioned mM1-expressing WT and Δ*sppA* strains to assess hyphal killing by human primary neutrophils isolated from three independent donors (Figure 6C and Supplemental Movies 1 and 2). In agreement with our *in vivo* data, human neutrophils consistently killed Δ*sppA* hyphae more effectively than the WT fungus (Figure 6C). These data support a critical role for SppA in the virulence of *A. fumigatus* during invasive infection by maintaining septal integrity to resist hyphal killing by both neutrophils and echinocandin antifungals.

## Discussion

We previously demonstrated that the *A. fumigatus* C_2_H_2_ transcription factor ZfpA promotes tolerance to FksA-targeting antifungals and contributes to virulence during invasive infection [18,19]. To identify downstream effectors, we performed untargeted proteomic analysis of WT, Δ*zfpA*, and OE::*zfpA A. fumigatus* 293 exposed to the echinocandin caspofungin. As a validation of our approach, we confirmed ZfpA-dependent induction of the oxylipin oxygenase PpoA, consistent with previous studies [18,20,21]. Among proteins regulated by ZfpA, the C2-domain protein encoded by AFUA_2G01790 showed the strongest caspofungin induced increase in both WT and OE::*zfpA* strains compared to Δ*zfpA*. We named this protein SppA (<u>s</u>eptal <u>p</u>ore protein) based on its recently characterized *A. oryzae* ortholog [22]. Proteomic and Northern blot analysis indicated that SppA induction during caspofungin stress is strongly dependent on ZfpA. Our characterization of deletion, complementation and overexpression *sppA* mutants establish SppA as a critical regulator of hyphal resilience and reveal an important link between septation and pathogenic fitness in *A. fumigatus*.

Functional septa are critical for tolerance to echinocandin induced cell wall damage in *A. fumigatus* [15,16,24]. By limiting cytoplasmic loss following apical tip lysis, septal pore plugging allows hyphae to compartmentalize segments, survive, and re-establish tip growth. Since *A. oryzae* SppA localizes to septa and contributes to septal integrity during hypotonic stress-induced tip lysis [22], we hypothesized that SppA would similarly promote resistance to FksA-targeting antifungals. Consistent with this paradigm, Δ*sppA* hyphae failed to effectively contain damage following lysis events induced by FksA targeting drugs and exhibited severe defects in septal morphology and function. These findings suggest that SppA promotes survival during antifungal stress by enabling the formation of septa capable of both withstanding mechanical stress and sealing damaged compartments.

The relationship between SppA and the C2-domain protein Inn1 in *Saccharomyces cerevisiae* may provide clues regarding its mechanism of action [25]. Inn1 couples membrane ingression, required for cytokinesis, with synthesis of the chitinous primary septum [26]. Although *Aspergillus* SppA proteins possess substantially extended C-terminal domains relative to Inn1 and appear to have diverged functionally from their yeast counterparts [22], their shared localization to sites of septation suggests that these proteins participate in evolutionarily related processes. The calcofluor-bright structures observed in SppA-overexpressing strains and constricting rings of SppA-GFP observed during septal formation are consistent with the possibility that SppA influences the localization or activity of chitin and cell wall synthetic machinery. However, direct evidence for such interactions is currently lacking and will require future biochemical investigation.

Beyond its role in antifungal tolerance, our data demonstrate that SppA is a major determinant of virulence. Deletion of *sppA* in two distinct *A. fumigatus* genetic backgrounds dramatically attenuated pathogenicity in two independent zebrafish models—hindbrain and swim bladder—indicating that the phenotype is robust and not strain specific. The observation that survival of Δ*sppA*-infected larvae approached that of PBS-injected controls highlights the importance of septal integrity during infection and identifies SppA as one of the most influential virulence determinants described for septum-associated proteins in *A. fumigatus*.

Our findings further suggest that SppA-mediated septal integrity contributes to fungal survival during neutrophil attack. While macrophages are involved in the clearance of ungerminated conidia, neutrophils are the principal phagocytes for controlling invasive growth [27]. Indeed, the virulence of Δ*sppA* was largely restored in the neutrophil impaired mpx:mCherry-2A-Rac2^D57N^ zebrafish line, further indicating that enhanced sensitivity to neutrophil-mediated damage contributes substantially to the observed virulence defect. However, the survival of neutrophil mutant larvae infected with Δ*sppA* was still significantly higher than those infected with WT or *sppA*+ suggesting that other factors may also contribute to the reduced virulence of this mutant. While *in vivo* treatment of neutrophil deficient larvae with micafungin did not impact survival of WT or complement strain infected fish, it completely abolished virulence of the Δ*sppA* mutant. We speculate that compromised septal architecture limits the ability of Δ*sppA* hyphae to compartmentalize and recover from localized host induced damage, analogous to their inability survive antifungal induced tip lysis. This conclusion was supported by *in vitro* microscopy-based assessment of primary human neutrophil mediated killing of WT and Δ*sppA A. fumigatus* hyphae wherein cytosolic fluorescent signal was completely lost in entire Δ*sppA* hyphae while the loss was restricted to individual compartments of WT hyphae allowing their regrowth and escape from neutrophil clusters. Thus, SppA may provide a common mechanism of protection against both host derived and antifungal drug induced hyphal injury.

Through use of double mutants, where SppA was overexpressed in a *ΔzfpA* background or deleted in an *OE::zfpA* background we are able to position SppA as a prominent component of hyphal integrity downstream of ZfpA. Moreover, the finding that overexpression of either gene in the absence of the other failed to restore resistance to FksA targeting drugs, suggests that SppA represents only one component of a broader ZfpA-regulated stress response. Additional ZfpA-dependent factors identified in our proteomic analysis likely cooperate with SppA in promoting survival during cell wall stress. Future work to define these downstream pathways will provide a more comprehensive understanding of how *A. fumigatus* coordinates adaptive responses to antifungal challenge.

In summary, we identify SppA as a previously unrecognized regulator of septal formation, antifungal tolerance, and virulence in *A. fumigatus*. Our data support a model in which ZfpA-induced SppA promotes proper septal formation necessary for effective septal pore sealing and thereby allowing hyphae to circumscribe cellular damage and preserve hyphal viability following antifungal or host mediated injury. Given that animals do not form septa and lack an orthologous C2-domain protein, SppA offers a potential target for the identification of novel antifungals to treat invasive fungal infections either in conjunction with current cell wall targeting drugs or as standalone agents.

## Materials and Methods

### Ethics statement

Animal care and use protocol M005405-A02 was approved by the University of Wisconsin - Madison Institutional Animal Care and Use Committee (IACUC). This protocol adheres to the federal Health Research Extension Act and the Public Health Service Policy on the Humane Care and Use of Laboratory Animals, overseen by the National Institutes of Health (NIH) Office of Laboratory Animal Welfare (OLAW).

### Fungal strains, media, and culture conditions

Wild type *A. fumigatus* Af293 and CEA10 are common clinical laboratory strains available through ATCC. All fungal strains used in this study are described in Table S1. Strains were maintained in 25-50% glycerol stock suspensions at -80 °C and grown on glucose minimal media (GMM) plates at 37 °C for three days. Conidia were collected into sterile water with 0.01% Tween 80. GMM plates and broth were prepared as previously described[28].

### Antifungals drugs

Caspofungin, micafungin, voriconazole, and amphotericin B were purchased from ApexBio. Enfumafungin was purchased from MedChemExpress. All antifungal drugs were dissolved at 10 mg/mL in DMSO and stored in 50 µL aliquots for single use at -20°C.

### Protein extraction and sample prep

Conidia of WT AF293, Δ*zfpA*, and OE::*zfpA* strains were inoculated at 5 x 10^7^ spores in 50ml of GMM and incubated at 37 °C and 250 RPM for 20 hours, followed by addition of either caspofungin (0.5ug/ml) or DMSO (0.025%) and incubation for an additional 4 hours. Each sample group consisted of four replicates. Tissue was collected into sterile miracloth, flash frozen, and lyophilized. Cells were broken into a powder by bead beating with Lysing Matrix Z beads (MP Biomedicals) at 4°C in a Retsch Mixer Mill MM400 for 5 cycles (60 seconds maximum frequency followed by 90 seconds rest). 1ml of 90% methanol was added to each sample to precipitate protein. Samples were then centrifuged at 4°C for 5 minutes at 14,000 x g and the supernatants removed. The protein pellets were resolubilized in lysis buffer (8 M urea, 10 mM TCEP, 40 mM CAA, 100 mM Tris) by 1 minute of bead beating. Each sample was diluted with 100mM Tris so the final concentration of urea was 1.5M, and protein concentration was determined by a BCA assay (Thermo Pierce). 100µg of protein was digested with Trypsin (Promega) and Lys-C (Sigma) in a 50:1 ratio of protein/enzyme overnight on a rocker at room temperature. Enzyme activity was quenched by adding trifluoroacetic acid (TFA) to a final pH below 2.0. The peptides were then desalted using 10mg Strata-X Polymeric Solid Phase Extraction cartridges (Phenomenex) and eluted in 300µl of elution buffer (80% ACN, 0.2% TFA). Samples were then dried with the SpeedVac Vacuum Concentrator before proteome analysis.

### LC-MS analysis

Peptide samples (1 µg per injection) were run on a Vanquish Neo UHPLC (Thermo Scientific) interfaced to an Orbitrap Ascend Tribrid mass spectrometer (Thermo Scientific) using a Nanospray Flex source. Chromatographic separation was performed on a 40 cm fused-silica column (75 µm inner diameter, 360 µm outer diameter; Polymicro Technologies) that was etched and packed in-house with 1.7 µm C18 particles (Waters). Peptides were separated on a Vanquish Neo nanoLC (Thermo Fisher) at 0.3 µL/min using a 73 min active gradient from 3% to 46% mobile phase B. Mobile phase A was 0.2% formic acid in water, and mobile phase B was 80% acetonitrile with 0.2% formic acid in water. MS data were acquired in positive-mode DDA on an Orbitrap Ascend Tribrid (Thermo Fisher). Full MS1 scans (Orbitrap) were collected every 1 s at 240,000 resolution over 300–1350 m/z (AGC target 1×10^6^; normalized AGC 250%; max injection time 50 ms; RF lens 30%). Monoisotopic peak determination was set to “peptide,” the isolation window center to “most abundant peak,” and precursors with charge states 2–5 were selected with dynamic exclusion of 20 s (±5 ppm). MS2 spectra were acquired in the ion trap with a 0.8 m/z isolation window using HCD at 24% normalized collision energy, Turbo scan rate, and a 150–1350 m/z scan range (normalized AGC 250%; max injection time 12 ms). The proteomics raw data is accessible on the MassIVE database with accession number MSV000102436.

### Comparative analysis of proteomic data

Raw LC–MS files were processed in MaxQuant (v2.6.1.0) using default parameters with label-free quantification (MaxLFQ), searching against the *A. fumigatus* Af293 reference proteome (FungiDB release 68). Downstream analyses used LFQ intensities that were TIC-normalized across runs and log2-transformed. Missing values were imputed by random sampling from a low-intensity distribution corresponding to the bottom 1% of observed values, constrained to be greater than the global minimum non-missing intensity. Log2 fold-changes were computed from group mean log2 intensities, and statistical significance was evaluated using two-sided independent-samples t-tests.

### Strain construction and Southern blotting

To generate gene deletion constructs of *A. fumigatus*, two 1 kb DNA fragments immediately upstream and downstream of each open reading frame (ORF), were amplified by PCR from WT Af293 genomic DNA, and were fused to a 2 kb *A. parasiticus pyrG* or *A. fumigatus argB* fragment from pJW24 or WT Af293 gDNA using double joint PCR [29,30]. To generate complemented strains two 1 kb DNA fragments immediately upstream and downstream of the *fcyB* ORF were amplified by PCR from WT Af293 genomic DNA and were fused to a 3.9 kb fragment containing the entire *sppA* locus including 1 kb upstream of the predicted ATG and 200 bp downstream of the predicted stop site. Transformants were grown on selective media containing 5-fluorocytosine (5-FC, 10 µg/ML) (Cayman Chemical Company) as previously published [31]. To generate overexpression constructs, two 1 kb fragments immediately upstream and downstream of the translational start site were amplified by PCR from *WT A. fumigatus* Af293 genomic DNA and *A.para.pyrG::A.nid.gpdA(p)* or *A.fu.argB*::*A. nidulans gpdA(p)* as the selectable marker and overexpression promoter were amplified from the plasmids pJMP9 or pJMP10 [32]. These three fragments were fused by double joint PCR. To create a *pyrG* auxotrophs of the OE::*zfpA* strains, the *pyrG* marker was removed by transforming TJW214.2 with a *pyrG* excision construct. The construct consisted of two 1 kb DNA fragments from the upstream and downstream regions of the *pyrG* marker at the OE::*zfpA* locus joined by PCR. Transformants were grown on GMM containing 5-fluoroorotic Acid (FOA, 1 mg/mL) (Thermo Scientific) and uracil (0.5 g/L)/uridine (0.5g/L). To create a *pyrG* auxotrophs of the OE::*sppA* strains, the *pyrG* marker was removed by transforming TJW480.1 with a *pyrG* excision construct. The construct consisted of two 1 kb DNA fragments from the upstream and downstream regions of the *pyrG* marker at the OE::*sppA* locus joined by PCR. Transformants were grown on selective media containing 5-fluoroorotic Acid (FOA, 1 mg/mL) and uracil (0.5 g/L)/uridine (0.5g/L). Parental strains were transformed following the previously described approach [33]. Single integration of the transformation construct was confirmed by Southern blotting using restriction enzyme digest with both dCTP-αP^32^ (Perkin Elmer) labeled 5′ and 3′ flanks of each construct.

### Germling lysis experiments

For each experimental condition, approximately 8,000 conidia were inoculated into three independent wells of a 48-well plate in 0.4 mL GMM broth with 0.01% w/v yeast extract. The plate was incubated on a Nikon Eclipse Ti Inverted Microscope in a heated microscope enclosure (OKO Labs, Burlingame, CA) at 37 °C for 4 hours before imaging. During this pre-incubation, three XY imaging positions were set in each well with at least 33 visible spores in each frame. Images were acquired every fifteen minutes at each XY position for 21 hours using a Nikon Plan Fluor 10X Ph1 DLL objective. Ninety-nine spores from the three frames in each well were selected blindly at time zero (4 hours post inoculation) and annotated in NIS-Elements AR software package (Version 5.30) to be assessed for lysis at 16 hours post inoculation. The number of annotated germlings lysed by 16 hours post inoculation was manually counted for each of the three wells per condition. Complete lysis of germlings was determined by the loss of normal diffraction of light compared to intact hyphae and confirmed by complete cessation of growth out to 24 hours as previously described. Statistical analyses were performed using GraphPad Prism (v11.0.0).

### Stress test dilution plating

To assess cell wall stress tolerance, 30 mL GMM square plates with the specified drug treatment or vehicle controls were inoculated with 10^5^, 10^4^, 10^3^, and 10^2^ spores of each strain and grown at 37 °C for 48 hours. All experiments were completed in triplicate with representative pictures presented in figures.

### Calcofluor white staining and epifluorescence imaging

Coverglass bottom 24-well plates were inoculated with 10^3^ conidia per mL of *A. fumigatus* in liquid GMM and incubated for 14 hours before staining with calcofluor white (Sigma-Aldrich) for epifluorescent microscopy using a Nikon Ti2E inverted microscope with Plan Apochromat Lambda D 20X air immersion and Plan Apochromat Lambda D 60X Oil immersion objectives. All image analysis was performed using ImageJ.

### Confocal imaging of SppA-GFP

Coverglass bottom 24-well plates were inoculated with 10^3^ conidia per mL of *A. fumigatus* in liquid GMM and incubated for 12-14 hours before live imaging on a Nikon Eclipse Ti2 microscope equipped with a Crest X-light V3531 spinning disc unit and a Kinetix22 monochrome camera. Imaging was performed using a Plan Apochromat 60X/1.42 NA objective.

### Zebrafish lines and maintenance

All zebrafish lines used in this study are listed in Supplemental Table 2. Adult zebrafish and larvae were maintained as previously described [34]. Larvae were anesthetized in E3 water (E3) + 0.2 mg/mL Tricaine (ethyl 3-aminobenzoate, Sigma) prior to all experiments.

### Larval zebrafish microinjection and survival studies

Larvae (2 dpf or 4 dpf respectively) were anesthetized and 3 nL of *A. fumigatus* conidial suspension was microinjected into the hindbrain ventricle via the otic vesicle or the developing swim bladder as previously described [34,35]. A conidial stock of 1.5e8 spores per mL was mixed 2:1 with 1% Phenol Red prior to injection to visualize the inoculum in the hindbrain. After injection larvae were rinsed 3X with E3 without methylene blue (E3-MB) to rinse off Tricaine and remained in E3-MB throughout all experiments. Larvae were transferred to individual wells of a 96-well plate for survival experiments or individual wells of 24- or 48-well plates for imaging experiments. For survival experiments, larvae were checked daily for 7 days and considered dead if there was no visible heartbeat. To determine the number of conidia injected for each experiment, 4 larvae/condition were collected after injection and individually added to microcentrifuge tubes in 90 μL 1X PBS with 500 μg/mL kanamycin and 500 μg/mL gentamycin. Larvae were homogenized for 15 sec using a mini-bead beater and plated on solid GMM plates. Colony forming units (CFUs) were counted after two-day incubation at 37°C to ensure similar average conidial inocula between strains (i.e. < 40% difference between strains within an experiment). Statistical analyses were performed using R (v4.5.1).

### Primary human neutrophil collection and isolation

All blood samples were obtained from healthy donors and were drawn according to the University of Wisconsin-Madison Minimal Risk Research Institutional Review Board-approved protocol (ID: 2017–0032) per the Declaration of Helsinki. Formal written consent was obtained from donors prior to blood draw. Neutrophils were isolated immediately after blood collection using the MACSxpress Whole Blood Neutrophil Isolation Kit (Miltyeni Biotec #130-104-434) and manufacturer instructions. Neutrophils were centrifuged for 5 min at 200 x g and the pellet was resuspended in 1 mL PBS for counting. Neutrophils were centrifuged again and resuspended to a final concentration of 10^6^ cells/mL in RPMI + 4% pooled human serum and used immediately.

### In vitro human neutrophil hyphal killing

A coverglass bottom 96-well plate was inoculated with 100 µL RPMI containing approximately 200 *A. fumigatus* conidia per well and incubated at 37°C and 5% CO_2_ for 12 hours. Following the 12-hour incubation, the media was replaced with 100 µL RPMI with 4% pooled human serum and 1 mg/mL Cultrex Ultimatrix (R&D Systems) and placed on a Nikon Eclipse Ti2 microscope equipped with a Tokai HIT stage top chamber set to 37°C and 5% CO_2_. Individual XY positions were preset on 50 hyphae of each strain for imaging. After one hour, 100 µL of RPMI containing 100,000 human primary neutrophils were added to each well. Images were acquired for 16 hours every 15 min using a CFI Super Fluor 20X objective. Hyphal viability was determined out to 12 hours by automatic thresholding in the far-red fluorescent channel using the RenyiEntropy formula in ImageJ v1.54. Statistical analysis was performed using GraphPad Prism (v11.0.0).

## Supporting information

Main Figures

## Acknowledgements

We thank the lab of Professor Mike Bromely in the Manchester Fungal Infection Group for generously providing the mMaroon1-expressing strain of *A. fumigatus* A1160.

## References

1. Denning DW. Global incidence and mortality of severe fungal disease. Lancet Infect Dis. 2024;24: e428–e438. doi:10.1016/s1473-3099(23)00692-8

2. Patterson TF, Thompson GR, Denning DW, Fishman JA, Hadley S, Herbrecht R, et al. Practice Guidelines for the Diagnosis and Management of Aspergillosis: 2016 Update by the Infectious Diseases Society of America. Clin Infect Dis. 2016;63: e1–e60. doi:10.1093/cid/ciw326

3. Barrett JP, Vardulaki KA, Conlon C, Cooke J, Daza-Ramirez P, Evans EGV, et al. A systematic review of the antifungal effectiveness and tolerability of amphotericin B formulations. Clin Ther. 2003;25: 1295–1320. doi:10.1016/s0149-2918(03)80125-x

4. Herbrecht R, Denning DW, Patterson TF, Bennett JE, Greene RE, Oestmann J-W, et al. Voriconazole versus Amphotericin B for Primary Therapy of Invasive Aspergillosis. N Engl J Med. 2002;347: 408–415. doi:10.1056/nejmoa020191

5. Gautam I, Ankur A, Singh K, Jacob A, Doering TL, Gow NAR, et al. Breaking down the wall: Solid-state NMR illuminates how fungi build and remodel diverse cell walls. PLOS Pathog. 2025;21: e1013678. doi:10.1371/journal.ppat.1013678

6. Gow NAR, Latge J-P, Munro CA. The Fungal Cell Wall: Structure, Biosynthesis, and Function. Microbiol Spectr. 2017;5: 10.1128/microbiolspec.funk-0035-2016. doi:10.1128/microbiolspec.funk-0035-2016

7. Szymański M, Chmielewska S, Czyżewska U, Malinowska M, Tylicki A. Echinocandins – structure, mechanism of action and use in antifungal therapy. J Enzym Inhib Med Chem. 2022;37: 876–894. doi:10.1080/14756366.2022.2050224

8. Kurtz MB, Heath IB, Marrinan J, Dreikorn S, Onishi J, Douglas C. Morphological effects of lipopeptides against Aspergillus fumigatus correlate with activities against (1,3)-beta-D-glucan synthase. Antimicrob Agents Chemother. 1994;38: 1480–1489. doi:10.1128/aac.38.7.1480

9. Mroczyńska M, Brillowska-Dąbrowska A. Review on Current Status of Echinocandins Use. Antibiotics. 2020;9: 227. doi:10.3390/antibiotics9050227

10. Bowman JC, Hicks PS, Kurtz MB, Rosen H, Schmatz DM, Liberator PA, et al. The Antifungal Echinocandin Caspofungin Acetate Kills Growing Cells of Aspergillus fumigatus In Vitro. Antimicrob Agents Chemother. 2002;46: 3001–3012. doi:10.1128/aac.46.9.3001-3012.2002

11. Ingham CJ, Schneeberger PM. Microcolony Imaging of Aspergillus fumigatus Treated with Echinocandins Reveals Both Fungistatic and Fungicidal Activities. PLoS ONE. 2012;7: e35478. doi:10.1371/journal.pone.0035478

12. Angulo DA, Alexander B, Rautemaa-Richardson R, Alastruey-Izquierdo A, Hoenigl M, Ibrahim AS, et al. Ibrexafungerp, a Novel Triterpenoid Antifungal in Development for the Treatment of Mold Infections. J Fungi. 2022;8: 1121. doi:10.3390/jof8111121

13. Lockhart SR, Chowdhary A, Gold JAW. The rapid emergence of antifungal-resistant human-pathogenic fungi. Nat Rev Microbiol. 2023;21: 818–832. doi:10.1038/s41579-023-00960-9

14. Moreno-Velásquez SD, Seidel C, Juvvadi PR, Steinbach WJ, Read ND. Caspofungin-Mediated Growth Inhibition and Paradoxical Growth in Aspergillus fumigatus Involve Fungicidal Hyphal Tip Lysis Coupled with Regenerative Intrahyphal Growth and Dynamic Changes in β-1,3-Glucan Synthase Localization. Antimicrob Agents Chemother. 2017;61: 10.1128/aac.00710-17. doi:10.1128/aac.00710-17

15. Souza ACO, Martin-Vicente A, Nywening AV, Ge W, Lowes DJ, Peters BM, et al. Loss of Septation Initiation Network (SIN) kinases blocks tissue invasion and unlocks echinocandin cidal activity against Aspergillus fumigatus. PLoS Pathog. 2021;17: e1009806. doi:10.1371/journal.ppat.1009806

16. Thorn HI, Guruceaga X, Martin-Vicente A, Nywening AV, Xie J, Ge W, et al. MOB-mediated regulation of septation initiation network (SIN) signaling is required for echinocandin-induced hyperseptation in Aspergillus fumigatus. mSphere. 2024;9: e00695–23. doi:10.1128/msphere.00695-23

17. Widanage MCD, Gautam I, Sarkar D, Mentink-Vigier F, Vermaas JV, Ding S-Y, et al. Adaptative survival of Aspergillus fumigatus to echinocandins arises from cell wall remodeling beyond β−1,3-glucan synthesis inhibition. Nat Commun. 2024;15: 6382. doi:10.1038/s41467-024-50799-8

18. Calise DG, Park SC, Bok JW, Goldman GH, Keller NP. An oxylipin signal confers protection against antifungal echinocandins in pathogenic aspergilli. Nat Commun. 2024;15: 3770. doi:10.1038/s41467-024-48231-2

19. Schoen TJ, Calise DG, Bok JW, Giese MA, Nwagwu CD, Zarnowski R, et al. Aspergillus fumigatus transcription factor ZfpA regulates hyphal development and alters susceptibility to antifungals and neutrophil killing during infection. PLOS Pathog. 2023;19: e1011152. doi:10.1371/journal.ppat.1011152

20. Calise DG, Michaelis ML, Park SC, Wagner AS, Keller NP. ZfpA-regulated chitin synthesis in Aspergillus fumigatus hyphae determines fungicidal tip lysis by FksA-targeting antifungals. Antimicrob Agents Chemother. 2026; e0176925. doi:10.1128/aac.01769-25

21. Niu M, Steffan BN, Fischer GJ, Venkatesh N, Raffa NL, Wettstein MA, et al. Fungal oxylipins direct programmed developmental switches in filamentous fungi. Nat Commun. 2020;11: 5158. doi:10.1038/s41467-020-18999-0

22. Mamun MdAA, Cao W, Nakamura S, Maruyama J. Large-scale identification of genes involved in septal pore plugging in multicellular fungi. Nat Commun. 2023;14: 1418. doi:10.1038/s41467-023-36925-y

23. Schoen TJ, Huttenlocher A, Keller NP. Guide to the Larval Zebrafish-Aspergillus Infection Model. Curr Protoc. 2021;1: e317. doi:10.1002/cpz1.317

24. Dichtl K, Samantaray S, Aimanianda V, Zhu Z, Prévost M, Latgé J, et al. Aspergillus fumigatus devoid of cell wall β-1,3-glucan is viable, massively sheds galactomannan and is killed by septum formation inhibitors. Mol Microbiol. 2015;95: 458–471. doi:10.1111/mmi.12877

25. Sanchez-Diaz A, Marchesi V, Murray S, Jones R, Pereira G, Edmondson R, et al. Inn1 couples contraction of the actomyosin ring to membrane ingression during cytokinesis in budding yeast. Nat Cell Biol. 2008;10: 395–406. doi:10.1038/ncb1701

26. Nishihama R, Schreiter JH, Onishi M, Vallen EA, Hanna J, Moravcevic K, et al. Role of Inn1 and its interactions with Hof1 and Cyk3 in promoting cleavage furrow and septum formation in S. cerevisiae. J Cell Biol. 2009;185: 995–1012. doi:10.1083/jcb.200903125

27. Mircescu MM, Lipuma L, Rooijen N van, Pamer EG, Hohl TM. Essential Role for Neutrophils but not Alveolar Macrophages at Early Time Points following Aspergillus fumigatus Infection. J Infect Dis. 2009;200: 647–656. doi:10.1086/600380

28. Shimizu K, Keller NP. Genetic Involvement of a cAMP-Dependent Protein Kinase in a G Protein Signaling Pathway Regulating Morphological and Chemical Transitions in Aspergillus nidulans. Genetics. 2001;157: 591–600. doi:10.1093/genetics/157.2.591

29. Bok JW, Soukup AA, Chadwick E, Chiang Y, Wang CCC, Keller NP. VeA and MvlA repression of the cryptic orsellinic acid gene cluster in Aspergillus nidulans involves histone 3 acetylation. Mol Microbiol. 2013;89: 963–974. doi:10.1111/mmi.12326

30. Calvo AM, Bok J, Brooks W, Keller NP. veA Is Required for Toxin and Sclerotial Production in Aspergillus parasiticus. Appl Environ Microbiol. 2004;70: 4733–4739. doi:10.1128/aem.70.8.4733-4739.2004

31. Storer ISR, Sastré-Velásquez LE, Easter T, Mertens B, Dallemulle A, Bottery M, et al. Shining a light on the impact of antifungals on Aspergillus fumigatus subcellular dynamics through fluorescence imaging. Antimicrob Agents Chemother. 2024;68: e00803–24. doi:10.1128/aac.00803-24

32. Soukup AA, Farnoodian M, Berthier E, Keller NP. NosA, a transcription factor important in Aspergillus fumigatus stress and developmental response, rescues the germination defect of a laeA deletion. Fungal Genet Biol. 2012;49: 857–865. doi:10.1016/j.fgb.2012.09.005

33. Bok JW, Keller NP. LaeA, a Regulator of Secondary Metabolism in Aspergillus spp. Eukaryot Cell. 2004;3: 527–535. doi:10.1128/ec.3.2.527-535.2004

34. Knox BP, Deng Q, Rood M, Eickhoff JC, Keller NP, Huttenlocher A. Distinct Innate Immune Phagocyte Responses to Aspergillus fumigatus Conidia and Hyphae in Zebrafish Larvae. Eukaryot Cell. 2014;13: 1266–1277. doi:10.1128/ec.00080-14

35. Gratacap RL, Bergeron AC, Wheeler RT. Modeling Mucosal Candidiasis in Larval Zebrafish by Swimbladder Injection. J Vis ExpL: JoVE. 2014; e52182–e52182. doi:10.3791/52182

