## Supplementary material for "The *Aspergillus fumigatus* C2-Domain Protein SppA is required for septal integrity and alters susceptibility to echinocandins and neutrophil killing during infection": Main Figures

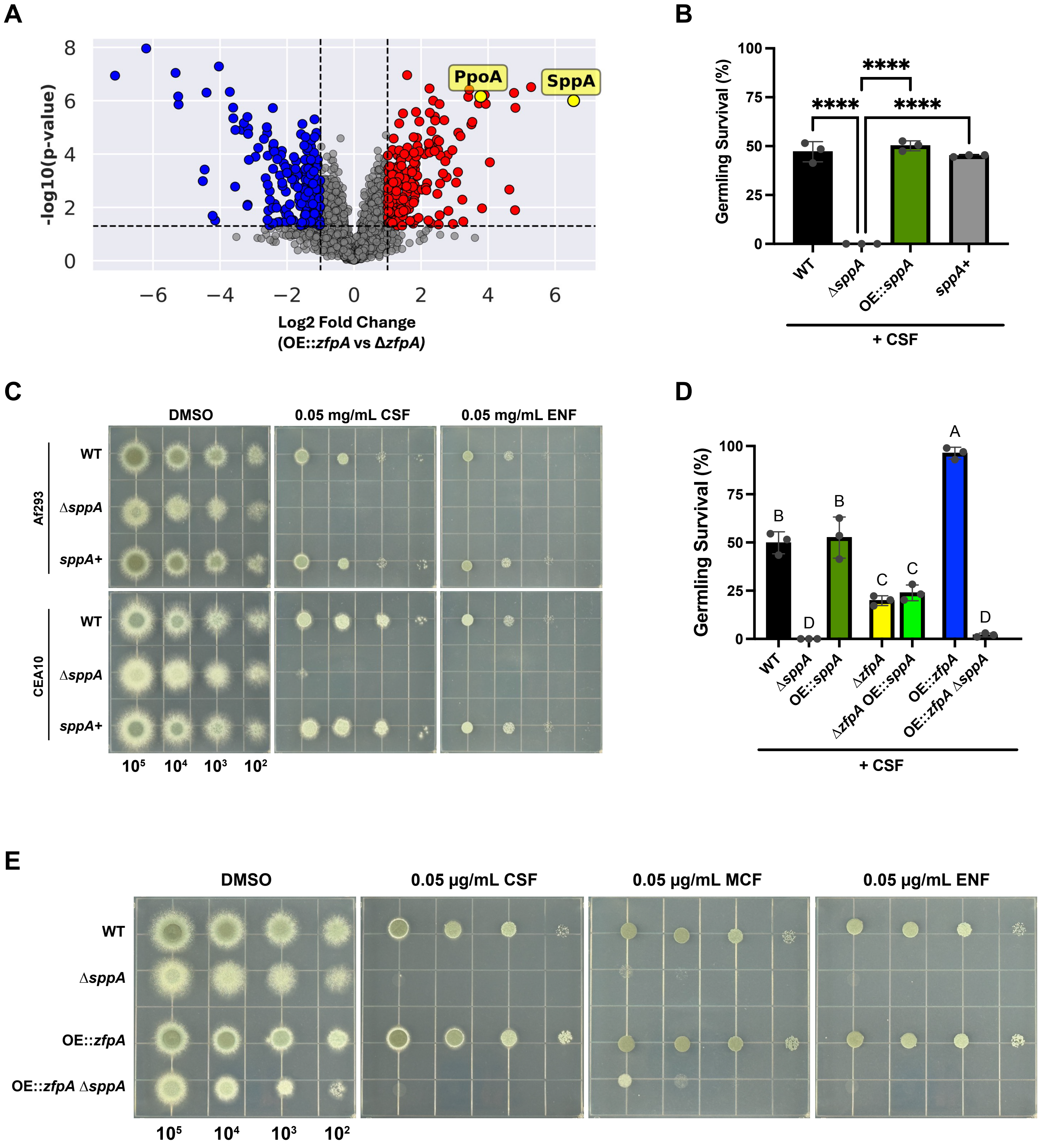
A
PpoA
SppA
-log10(p-value)
Log2 Fold Change
(OE::zfpA vs ΔzfpA)
B
Germling Survival (%)
****
****
****
WT
ΔsppA
OE::sppA
sppA+
+ CSF
C
DMSO
0.05 mg/mL CSF
0.05 mg/mL ENF
Af293
CEA10
WT
ΔsppA
sppA+
10⁵ 10⁴ 10³ 10²
D
Germling Survival (%)
A
B
B
C
C
D
D
WT
ΔsppA
OE::sppA
ΔzfpA
ΔzfpA OE::sppA
OE::zfpA
OE::zfpA ΔsppA
+ CSF
E
DMSO
0.05 μg/mL CSF
0.05 μg/mL MCF
0.05 μg/mL ENF
WT
ΔsppA
OE::zfpA
OE::zfpA ΔsppA
10⁵ 10⁴ 10³ 10²

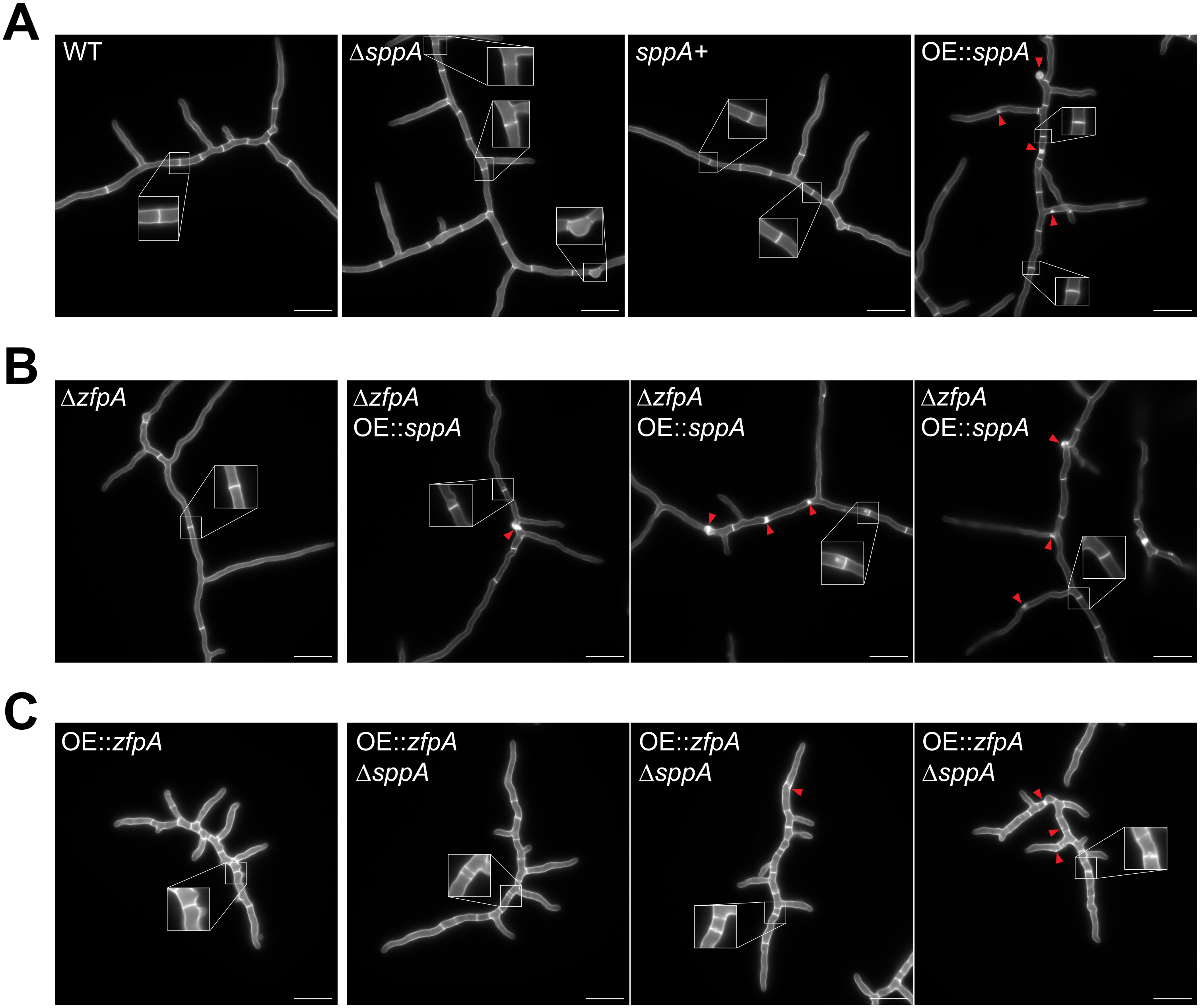
A
WT
ΔsppA
sppA+
OE::sppA
B
ΔzfpA
ΔzfpA OE::sppA
ΔzfpA OE::sppA
ΔzfpA OE::sppA
C
OE::zfpA
OE::zfpA ΔsppA
OE::zfpA ΔsppA
OE::zfpA ΔsppA

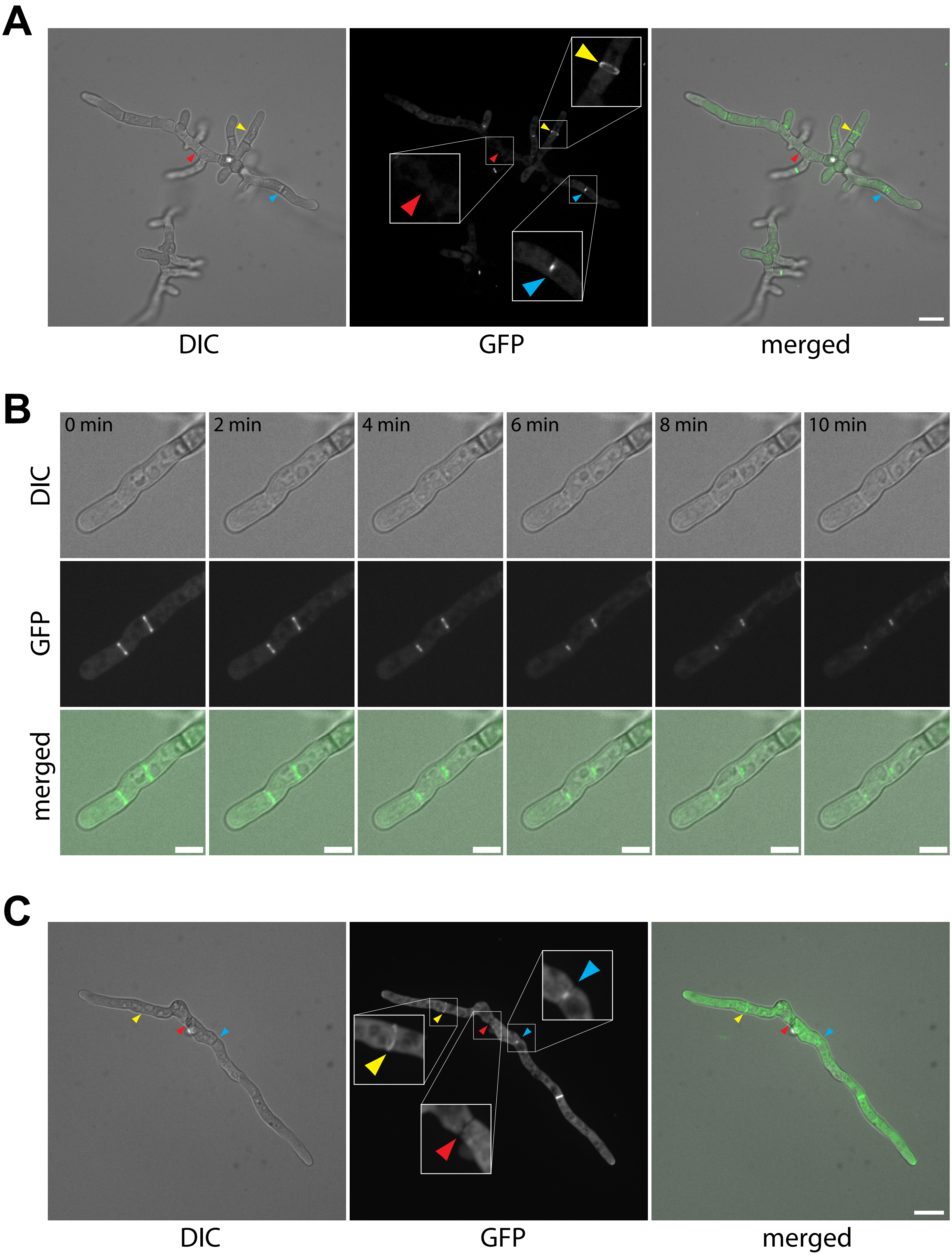
A
DIC
GFP
merged
B
0 min
2 min
4 min
6 min
8 min
10 min
DIC
GFP
merged
C
DIC
GFP
merged

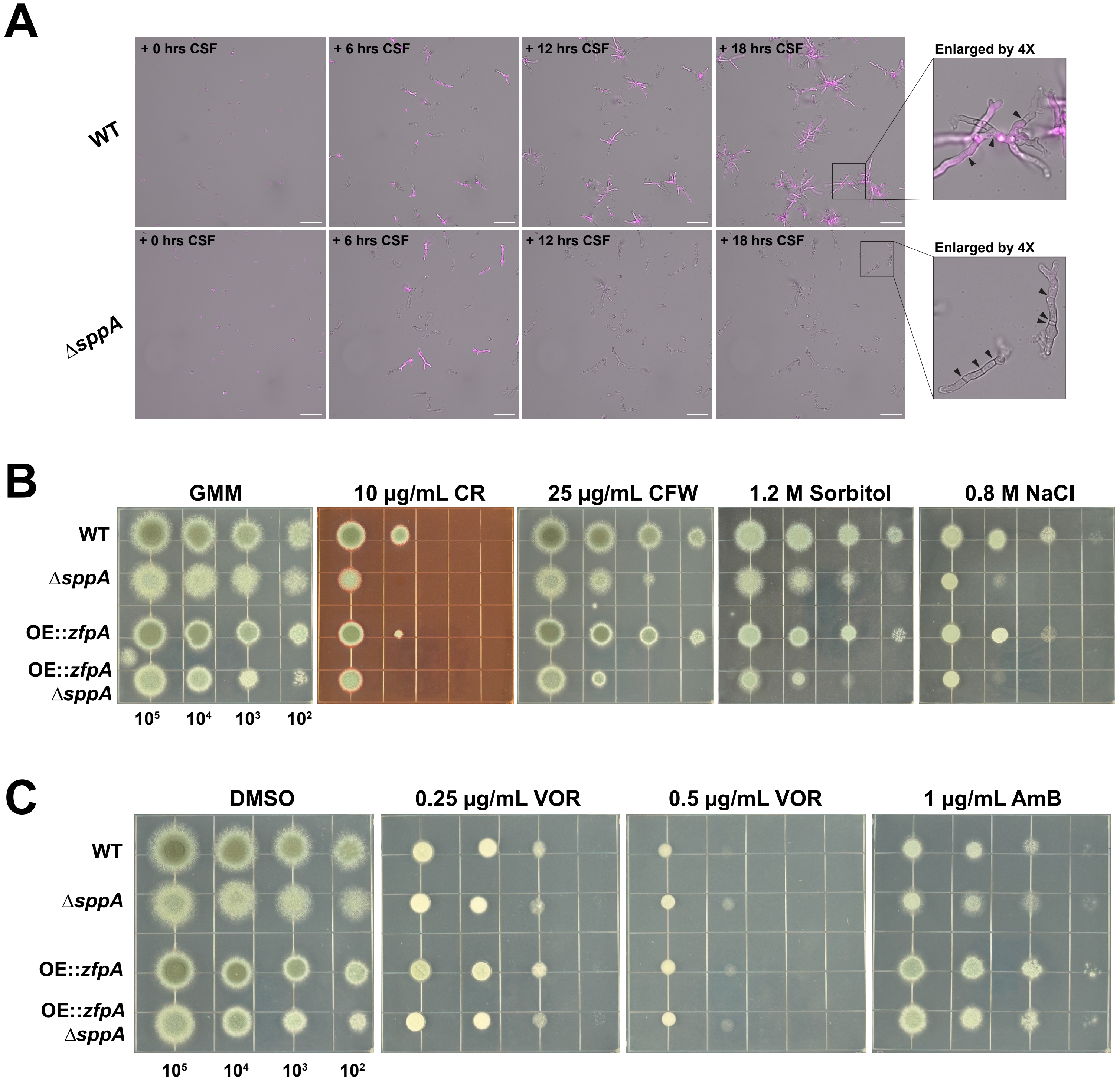
A
+ 0 hrs CSF
+ 6 hrs CSF
+ 12 hrs CSF
+ 18 hrs CSF
Enlarged by 4X
WT
ΔsppA
B
GMM
10 μg/mL CR
25 μg/mL CFW
1.2 M Sorbitol
0.8 M NaCl
WT
ΔsppA
OE::zfpA
OE::zfpA ΔsppA
10⁵ 10⁴ 10³ 10²
C
DMSO
0.25 μg/mL VOR
0.5 μg/mL VOR
1 μg/mL AmB
WT
ΔsppA
OE::zfpA
OE::zfpA ΔsppA
10⁵ 10⁴ 10³ 10²

A

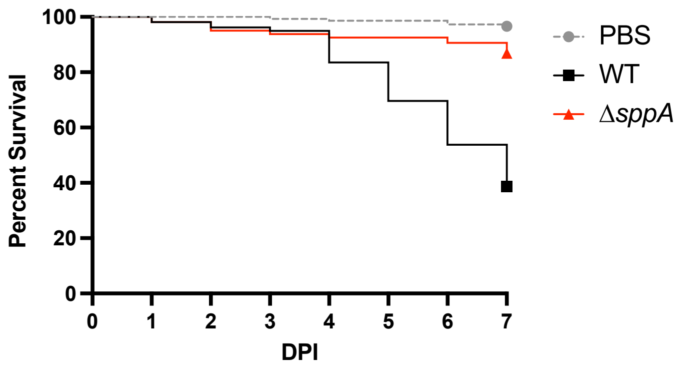


| Comparison | Hazard Ratio | P value |
|---|---|---|
| WT vs. PBS | 28.49 | < 0.0001 |
| Δ*sppA* vs PBS | 4.22 | 0.0114 |
| Δ*sppA* vs. WT | 0.1482 | < 0.0001 |

B

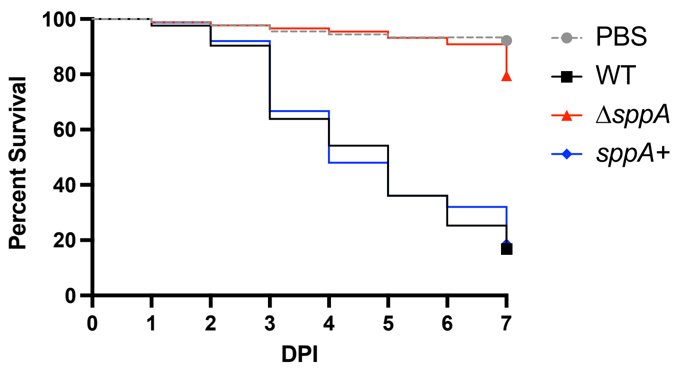


| Comparison | Hazard Ratio | P value |
|---|---|---|
| WT vs PBS | 12.55 | < 0.0001 |
| *sppA*+ vs PBS | 12.56 | < 0.0001 |
| Δ*sppA* vs WT | 0.1601 | < 0.0001 |
| *sppA*+ vs Δ*sppA* | 6.258 | < 0.0001 |

C

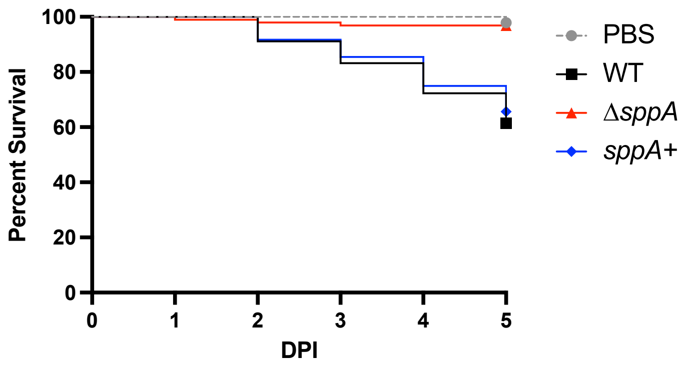


| Comparison | Hazard Ratio | P value |
|---|---|---|
| WT vs PBS | 22.62 | 0.0001 |
| *sppA*+ vs PBS | 19.53 | 0.0003 |
| Δ*sppA* vs WT | 0.0658 | < 0.0001 |
| *sppA*+ vs Δ*sppA* | 13.12 | 0.0001 |

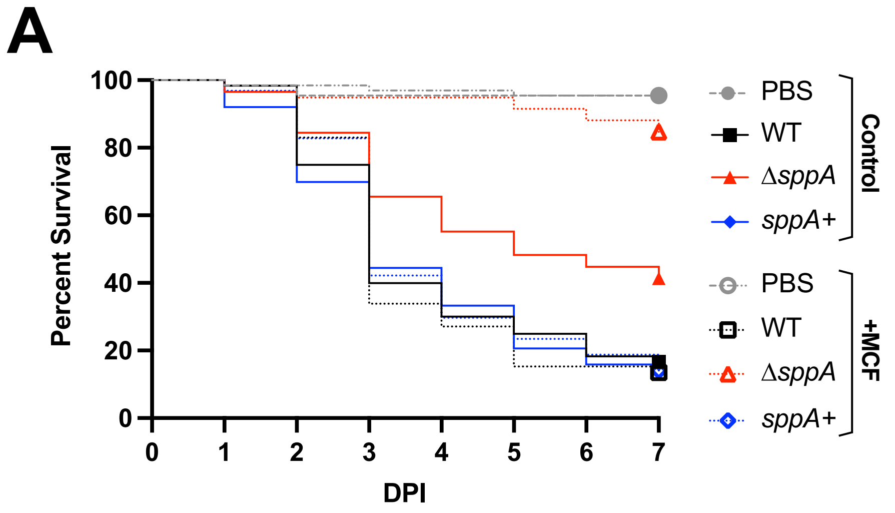


| Comparison | Hazard Ratio | P value |
|---|---|---|
| WT vs PBS | 39.68 | < 0.0001 |
| Δ*sppA* vs PBS | 20.70 | < 0.0001 |
| *sppA*+ vs PBS | 41.38 | < 0.0001 |
| Δ*sppA* vs WT | 0.5222 | 0.0219 |
| *sppA*+ vs Δ*sppA* | 1.999 | 0.0114 |
| WT (MCF) vs PBS (MCF) | 49.26 | < 0.0001 |
| Δ*sppA* (MCF) vs PBS (MCF) | n.s. | n.s. |
| *sppA*+ (MCF) vs PBS (MCF) | 44.52 | < 0.0001 |
| Δ*sppA* (MCF) vs WT (MCF) | 0.0735 | < 0.0001 |
| *sppA*+ (MCF) vs Δ*sppA* (MCF) | 12.30 | < 0.0001 |
| Δ*sppA* (MCF) vs Δ*sppA* | 0.1701 | < 0.0001 |

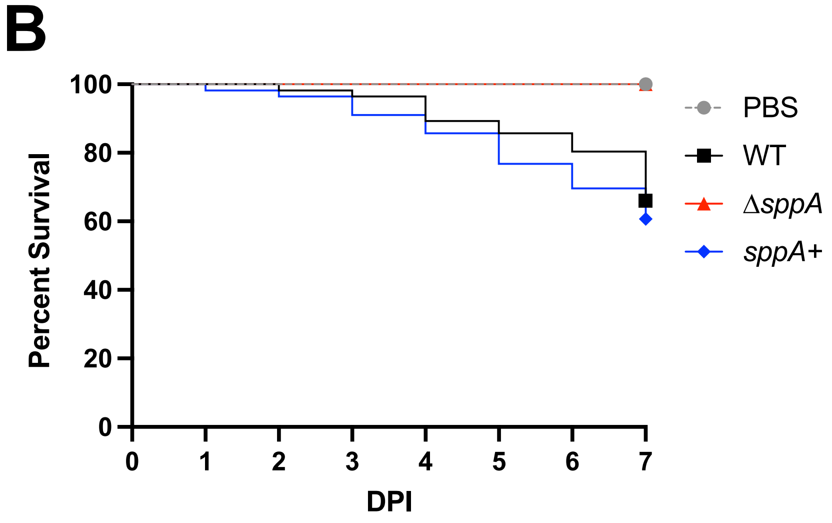


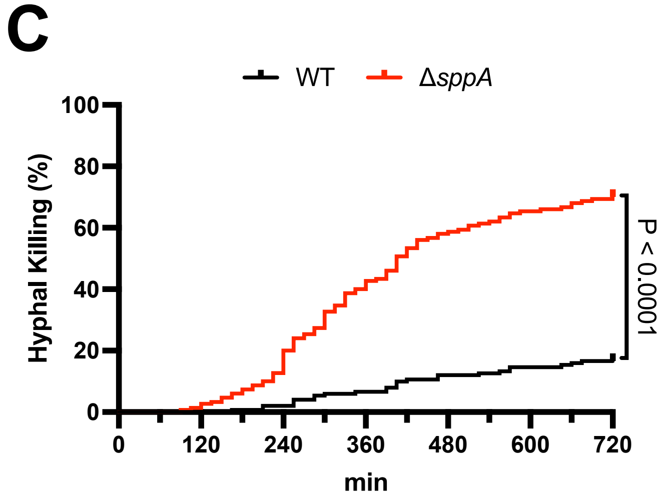


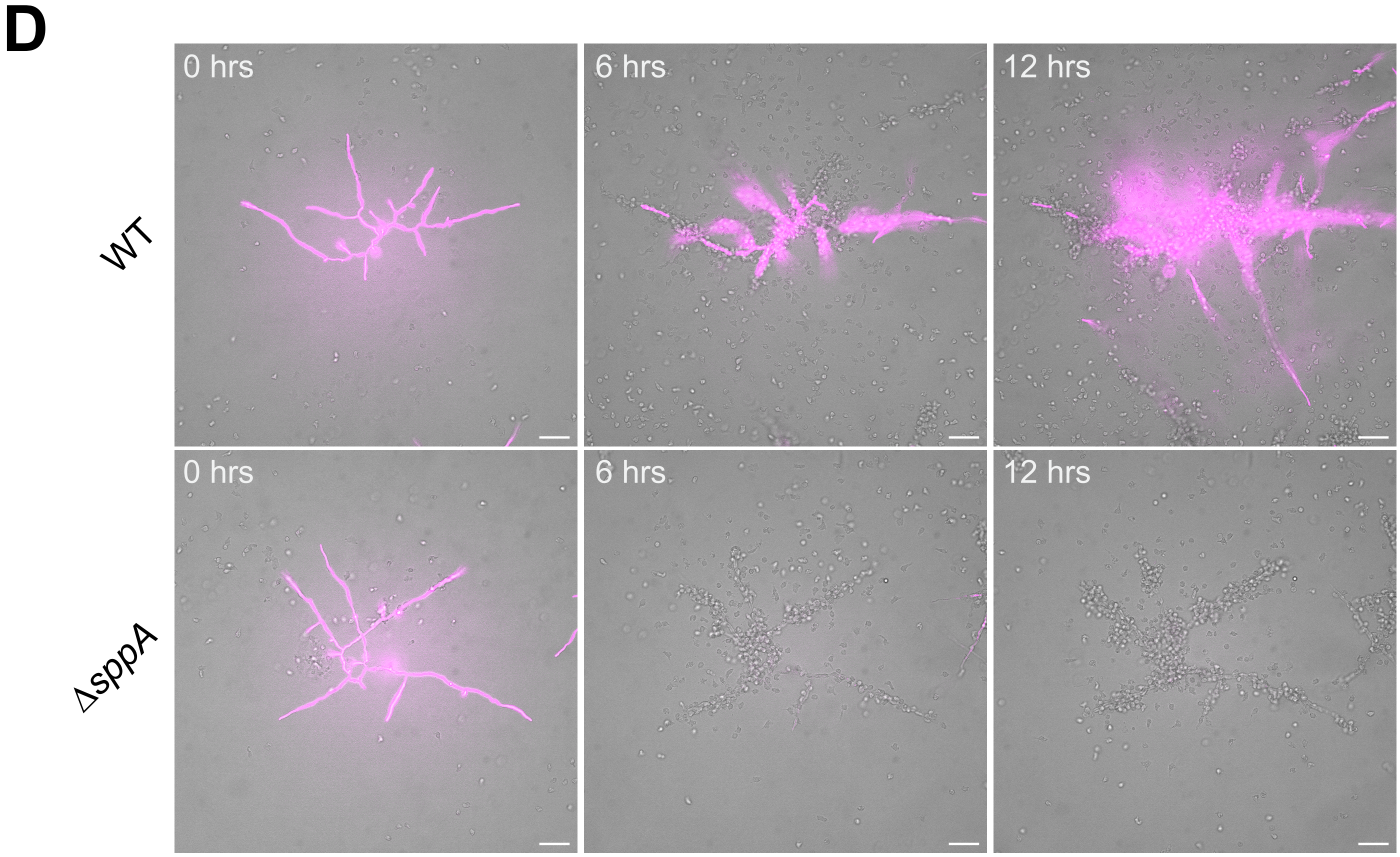
