## Supplemental Information for "The *Aspergillus fumigatus* C2-Domain Protein SppA is required for septal integrity and alters susceptibility to echinocandins and neutrophil killing during infection"

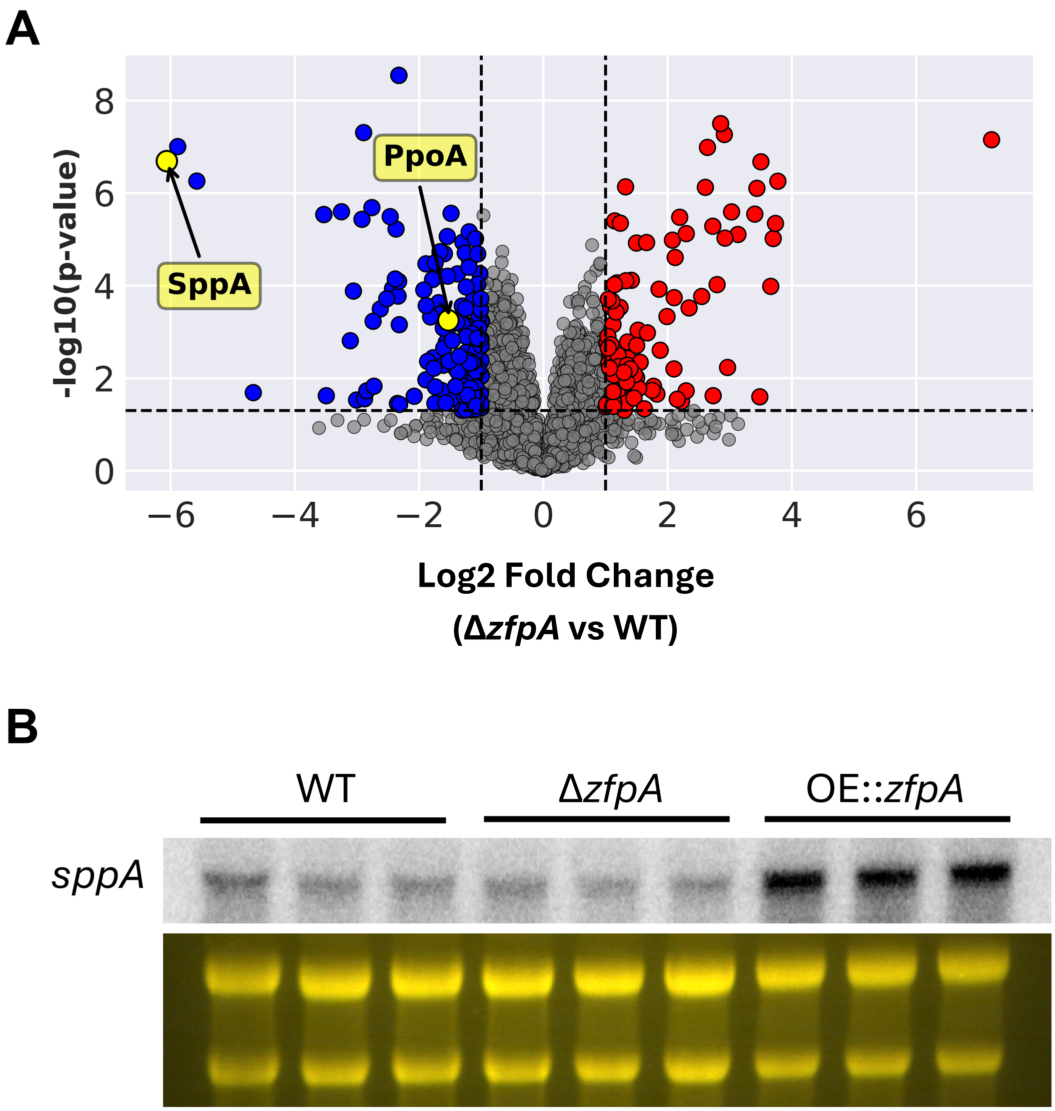


**Supplemental Figure 1: (A)** $Log_2FC$ abundance of *A. fumigatus* Af293 proteins in OE::*zfpA* relative to Δ*zfpA* extracted from mycelia after treatment with 0.5 µg/mL caspofungin (CSF) for four hours. Points representing SppA and PpoA are colored yellow. SppA was identified as the most downregulated protein. **(B)** Northern blot analysis of *sppA* expression in *A. fumigatus* Af293 after 24 hours in GMM at 37˚C and 250 RPM.

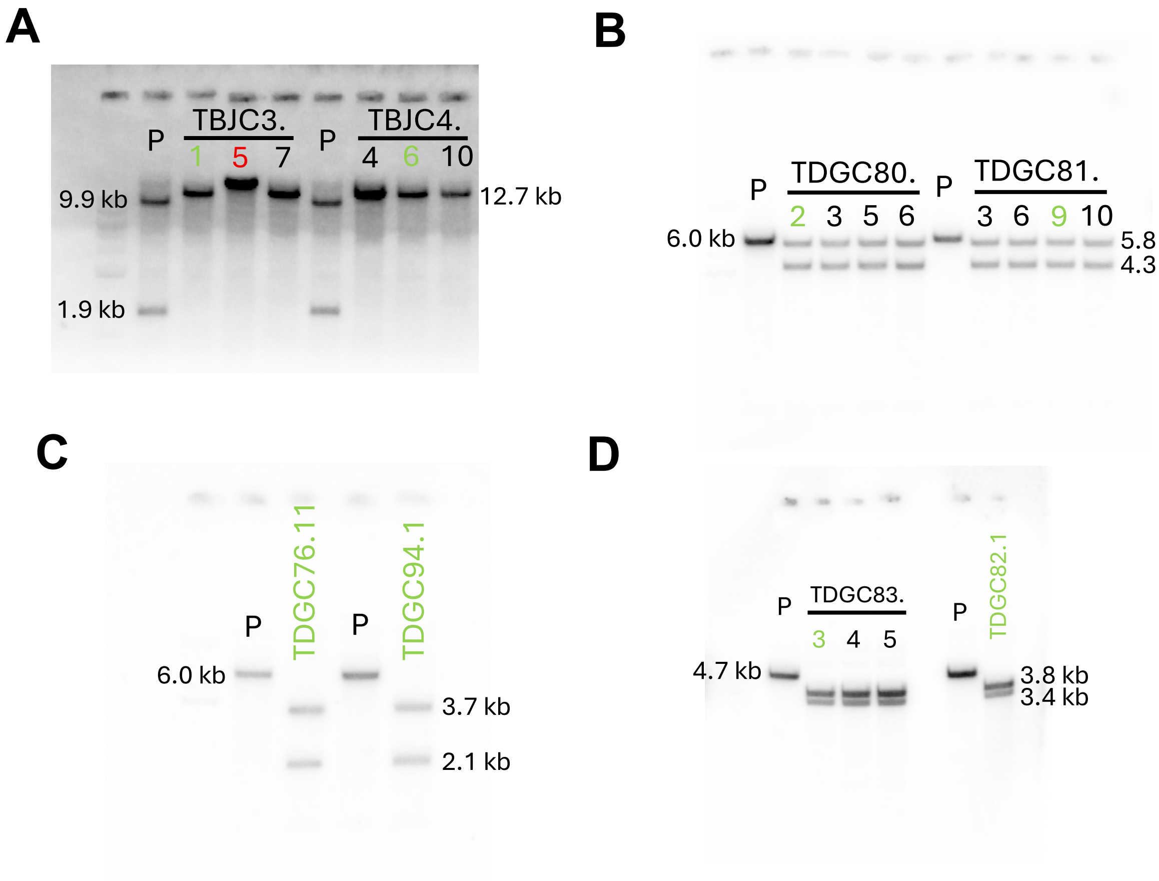


**Supplemental Figure 2: (A)** Southern blot of StuI digested genomic DNA of parental strains TDGC19.1 and TDGC12.2 and the resulting Δ*sppA* mutants. **(B)** Southern blot of PvuII-HF digested genomic DNA of parental strains TDGC19.1 and TDGC11.3 and the resulting OE::*sppA* mutants. **(C)** Southern blot of StuI digested genomic DNA of parental strains CEA17 *pyrG-* and TDGC90.1 and the resulting Δ*sppA* mutants. **(D)** Southern blot of StuI digested genomic DNA of parental strains TBJC3.1 and TDGC76.11 and the resulting Δ*fcyB*::*sppA* mutants.

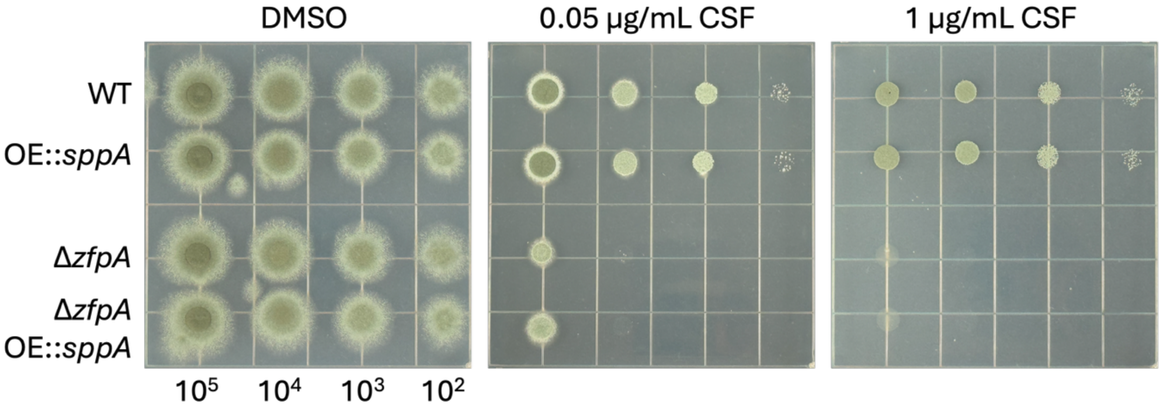


**Supplemental Figure 3:** *A. fumigatus* Af293 conidia spotted on GMM after incubation at 37˚C for 48 hours.

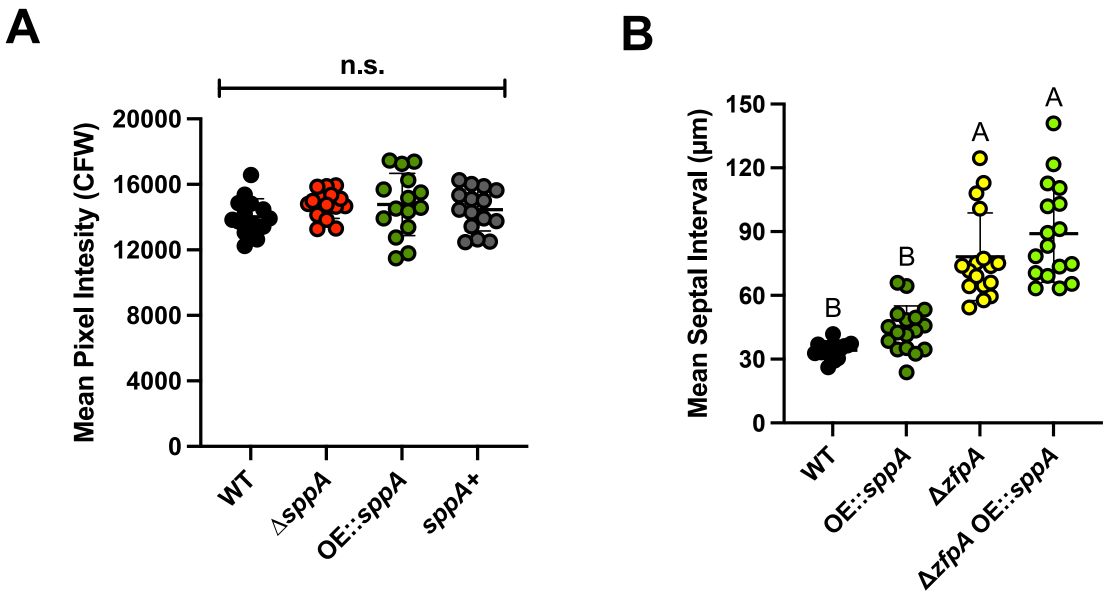


**Supplemental Figure 4: (A)** Mean CFW intensity per pixel of ten hyphae grown for 15 hours in GMM before staining and epifluorescence imaging. Points represent individual hyphae grown for sixteen hours in GMM. **(B)** Mean distance between apparently normal septa measured manually using ImageJ after by CFW staining and visualization by epifluorescence microscopy. Points represent individual hyphae grown for sixteen hours in GMM. Conditions with p values less than 0.05 calculated by one-way ANOVA with Tukey's multiple comparisons are indicated by distinct letters.

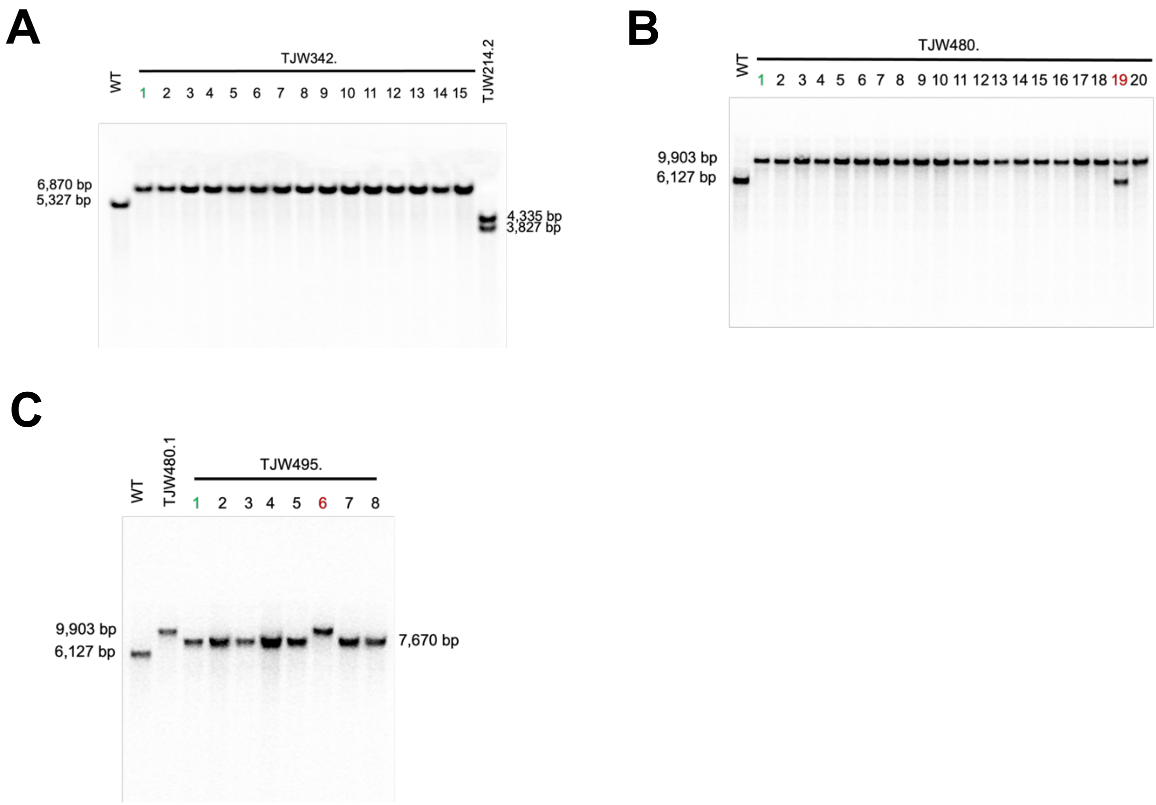


**Supplemental Figure 5: (A)** Southern blot of EcoRV digested genomic DNA of parental strain TJW214.2 and the resulting *pyrG*- mutants. **(B)** Southern blot of PvuI digested genomic DNA of parental strain TFYL80.1 and the resulting OE::*sppA* mutants. **(C)** Southern blot of PvuI digested genomic DNA of parental strain TJW480.1 and the resulting *pyrG*- mutants.

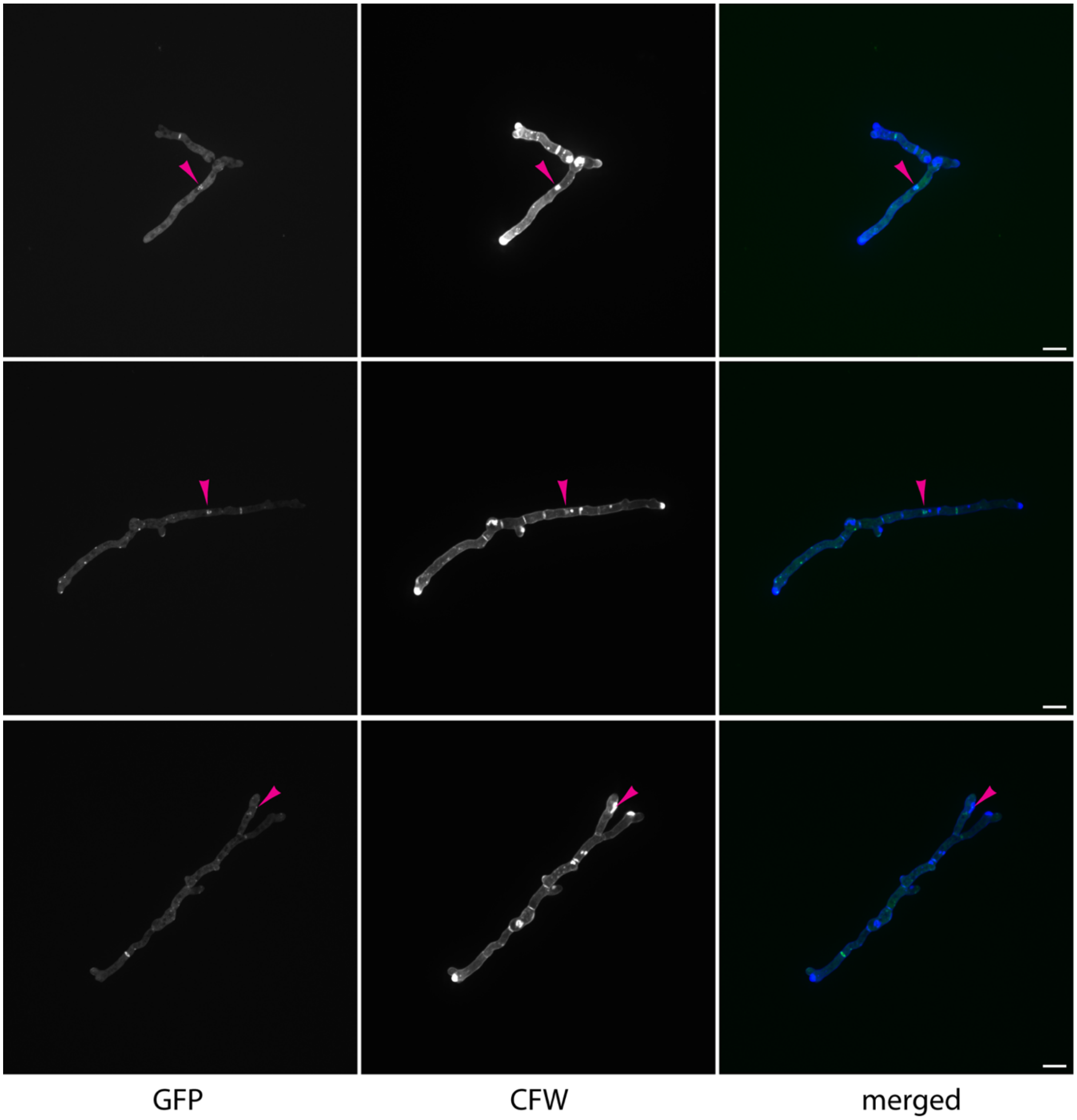


**Supplemental Figure 6:** Representative maximum intensity projections of Z-stack images collected every 0.2 µm by confocal fluorescence microscopy of OE::*sppA-GFP* hyphae grown for 13 hours in glucose minimal media before staining with 0.1 mg/mL calcofluor white (CFW). Magenta arrowheads denote colocalization of SppA-GFP puncta with brightly stained chitin patches in the cell wall. Scale bar represents 10 µm.

| STRAIN NAME | BACKGROUND | GENOTYPE | SOURCE |
|---|---|---|---|
| **TFYL81.5** | Af293 | Δ*akuA* | [1] |
| **TJW213.1** | Af293 | Δ*akuA*; Δ*zfpA* | [2] |
| **TJW214.2** | Af293 | Δ*akuA*; OE::*zfpA* | [2] |
| **TDGC19.1** | Af293 | Δ*akuA*; *argB1* | [3] |
| **TBJC3.1** | Af293 | Δ*akuA*; Δ*sppA*::*A.fu.argB* | This study |
| **TDGC80.2** | Af293 | Δ*akuA*; *A.fu.argB*::*A.n.gpdA*(p)::*sppA* | This study |
| **TDGC83.3** | Af293 | Δ*akuA*; Δ*sppA*::*A.fu.argB;* Δ*fcyB::sppA+* | This study |
| **A1160+** | CEA10 | Δ*akuB* | |
| **A1160-** | CEA10 | Δ*akuB*; *pyrG-* | |
| **TDGC76.11** | CEA10 | Δ*akuB*; Δ*sppA*::*A.p.pyrG* | This study |
| **TDGC82.1** | CEA10 | Δ*akuA*; Δ*sppA*::*A.p.pyrG;* Δ*fcyB::sppA+* | This study |
| **TDGC11.3** | Af293 | Δ*akuA*; *argB1;* Δ*zfpA::A.p.pyrG* | [3] |
| **TDGC12.2** | Af293 | Δ*akuA*; *argB1*; *A.p.pyrG*::*A.n.gpdA*(p)::*zfpA* | [3] |
| **TDGC81.9** | Af293 | Δ*akuA*; Δ*zfpA::A.p.pyrG;* *A.fu.argB*::*A.n.gpdA*(p)::sppA | This study |
| **TBJC4.4** | Af293 | Δ*akuA*; *A.p.pyrG*::*A.n.gpdA*(p)::*zfpA;* Δ*sppA::A.fu.argB* | This study |
| **MMAROON1** | CEA10 | Δ*akuB*; Δ*fcyB*::*A.n.gpdA*(p)::mMaroon1 | [4] |
| **TDGC90.1** | CEA10 | Δ*akuB*; Δ*fcyB*::*A.n.gpdA*(p)::mMaroon1; *pyrG-* | This study |
| **TDGC94.1** | CEA10 | Δ*akuB*; Δ*fcyB*::*A.n.gpdA*(p)::mMaroon1; Δ*sppA*::*A.p.pyrG* | This study |
| **TFYL80.1** | Af293 | Δ*akuA*; *pyrG1* | [5] |
| **TJW342.1** | Af293 | Δ*akuA*; *A.n.gpdA*(p)::*zfpA; pyrG-* | This study |
| **TJW480.1** | Af293 | Δ*akuA*; *A.p.pyrG*::*A.n.gpdA*(p)::*sppA* | This study |
| **TJW495.1** | Af293 | Δ*akuA*; *A.n.gpdA*(p)::*sppA; pyrG-* | This study |
| **TMLM9.1** | Af293 | Δ*akuA*; Δ*sppA::sppA-GFP-A.fu.pyrG* | This study |
| **TMLM10.1** | Af293 | Δ*akuA*; *A.n.gpdA*(p)::*sppA-GFP-A.fu.pyrG* | This study |
| **TMLM11.1** | Af293 | Δ*akuA*; *A.n.gpdA*(p)::*zfpA* Δ*sppA::sppA-GFP-A.fu.pyrG* | This study |

**Supplemental Table 1.** List of *A. fumigatus* strains used in this study.

| **Zebrafish (*Danio rerio*) Line** | **Source** |
|---|---|
| WT AB | ZIRC |
| Tg(mpx:mCherry-2A-Rac2$^{D57N}$) | [6] |
| Tg(mpx:mCherry-2A-Rac2$^{WT}$) | [6] |

**Supplemental Table 2.** List of *Danio rerio* strains used in this study.

| Name | Sequence |
|---|---|
| sppA_NorthernF | CCCGACTACTTCAGACTCAAGG |
| sppA_NorthernR | AGTATTGAGCAGCTTCGTGTCC |
| BC1 (KOsppA) | GTTTGACGATAGCGGAGCCG |
| BC2 (KOsppA) | aaaatttgtcttggatgcagaccgcgttcAGTGTAGTGTCCTGCGACTG |
| BC3 (KOsppA) | gatcaaatggatgtatgggtctctccttcGCGTAGCTTCTTCGAGGAGG |
| BC4 (KOsppA) | AACCCAAATCAACCAACGCC |
| A.f.argBF | gaaggagagacccatacatcc |
| A.f.argBR | agcttgaagtattatgggatgatg |
| sppA_5'flank_overhang R | CGATATCAAGCTTATCGATACCGTCGACGATGGCACCTTTCTC TCTGAAAGAGC |
| sppA_3'flank_overhang F | CGCTGCAGCCTCTCCGATTGTCGAATCGGCCAAAGTACCTATT GCACC |
| sppA_1kb3'flankR | GAGGAAAGCCTTTCCATCTCCC |
| sppA_5'flank_pJMP9ov erhangR | gtcctctcgggccatctgttcgtataagctGATGGCACCTTTCTCTCTGAAAG AGC |
| sppA_pJMP9overhang F | gctaccccgcttgagcagacatcaccATGGCTTCCAAACTACCCAGGATA GC |
| sppA_1kbORFR | TCTCGATGAAAGTCGTCAGGCG |
| sppA_confF | CCAATATTAACCATGCCGCCGG |
| sppA_confR | TCAGATCGATCCAGGTTTCGCC |
| fcyB_1kb5'flankF2 | TCTTTCCCTAGCGTCGGTGATG |
| fcyB_1kb3'flankR2 | TTGTTCGTCGTGACTTGTTGCG |
| fcyB_5'flank_sppA5'ove rhangR | gggacgagtctaacctgggtatctccAGTCAGTTTTGGACATTGGCCC |
| fcyB_5'flank_sppA3'ove rhangF | gcagaatgggggatcaataacgccccaaacGCTCTTGATGATAGAAGTGTG CGG |
| sppA_1kb5'flankF2 | ggagatacccaggttagactcgtcc |
| sppA_200bp3'flankR | gtttggggcgttattgatcccc |
| sppAc_confR | atgagttctccacagggatcgc |
| zfpAOE5F | TGACCATGATCTCCACTTCCCC |
| OEzfpApyrGrecyc5R | CTCATGTTTGACAGCTTATCATCGATAAGCGCAGACGTCCTAAG CTCGATAGTCGACTG |
| AppyrG recyF | GCTTATCGATGATAAGCTGTCAAAC |
| AppyrG recyR | GCTCTACCTACTTCGGAGAAGG |
| sppAOE5F | TCCTTTTGCTCGTCATGTCCCC |
| sppAOE5R | CCAATTCGCCCTATAGTGAGTCGTATTACGGATGGCACCTTTCT CTCTGAAAGAGCG |
| OEPyGF | CGTAATACGACTCACTATAGGGC |
| OEPyGR | GGTGATGTCTGCTCAAGCGGG |
| sppAOE3F | CAGCTACCCCGCTTGAGCAGACATCACCATGGCTTCCAAACTA CCCAGGATAGCC |
| sppAOE3R | GGATAGCTCTCTTCTTGTGGGG |
| OEsppApyrGrecyc5R | CTCATGTTTGACAGCTTATCATCGATAAGCGATGGCACCTTTCT CTCTGAAAGAGCG |
| 5F_5fl_out_GFP-sppA | CTCAATACTCCCCTCATTCGTCG |
| 5F_5fl_inn_GFP-sppA | GACGAAGAAGAGGAGGGACCTC |
| 3R_5fl_OH_GFP-sppA | CACCGGCTCCAGCGCCTGCACCAGCTCCcatagcgtacgttatcgcctt g |

| | |
|---|---|
| 5F_3fl_OH_GFP-sppA | CATCAGTGCCTCCTCTCAGACAGttatgagacctatgggtataacg |
| 3R_3fl_inn_GFP-sppA | CCTGCTCTTTACTTTTGCCGGTG |
| 3R_3fl_out_GFP-sppA | GCATCCCATCTACTGGATCACC |
| 5F_pFN03_GFP-sppA | GGAGCTGGTGCAGGCGCTG |
| 3R_gfp-pyrG_SppA | CTGTCTGAGAGGAGGCACTGATG |
| 5F_sequencing_sppA | GCCAATCCATCATCCAATCTGGTC |
| 3R_Afum_pyrG_conf | GTGCCAATCTGGATGGAGACAG |
| 3R_conf_GFP-sppA | GTCTTGTAGTTCCCGTCGTCTTTG |

**Supplemental Table 3.** List of oligonucleotides used in this study.

**Supplemental References**


1. Throckmorton K, Lim FY, Kontoyiannis DP, Zheng W, Keller NP. Redundant synthesis of a conidial polyketide by two distinct secondary metabolite clusters in Aspergillus fumigatus. Environ Microbiol. 2016;18: 246–259. doi:10.1111/1462-2920.13007

2. Schoen TJ, Calise DG, Bok JW, Giese MA, Nwagwu CD, Zarnowski R, et al. Aspergillus fumigatus transcription factor ZfpA regulates hyphal development and alters susceptibility to antifungals and neutrophil killing during infection. PLOS Pathog. 2023;19: e1011152. doi:10.1371/journal.ppat.1011152

3. Calise DG, Michaelis ML, Park SC, Wagner AS, Keller NP. ZfpA-regulated chitin synthesis in Aspergillus fumigatus hyphae determines fungicidal tip lysis by FksA-targeting antifungals. Antimicrob Agents Chemother. 2026; e0176925. doi:10.1128/aac.01769-25

4. Storer ISR, Sastré-Velásquez LE, Easter T, Mertens B, Dallemulle A, Bottery M, et al. Shining a light on the impact of antifungals on Aspergillus fumigatus subcellular dynamics through fluorescence imaging. Antimicrob Agents Chemother. 2024;68: e00803-24. doi:10.1128/aac.00803-24

5. Lim FY, Won TH, Raffa N, Baccile JA, Wisecaver J, Rokas A, et al. Fungal Isocyanide Synthases and Xanthocillin Biosynthesis in Aspergillus fumigatus. mBio. 2018;9: 10.1128/mbio.00785-18. doi:10.1128/mbio.00785-18

6. Deng Q, Yoo SK, Cavnar PJ, Green JM, Huttenlocher A. Dual Roles for Rac2 in Neutrophil Motility and Active Retention in Zebrafish Hematopoietic Tissue. Dev Cell. 2011;21: 735–745. doi:10.1016/j.devcel.2011.07.013
